# Single cell transcriptomics of primate sensory neurons identifies cell types associated with human chronic pain

**DOI:** 10.1101/2020.12.07.414193

**Authors:** Jussi Kupari, Dmitry Usoskin, Daohua Lou, Marc Parisien, Yizhou Hu, Michael Fatt, Peter Lönnerberg, Mats Spångberg, Bengt Eriksson, Nikolaos Barkas, Peter V Kharchenko, Karin Loré, Samar Khoury, Luda Diatchenko, Patrik Ernfors

## Abstract

Distinct types of dorsal root ganglion sensory neurons may have unique contributions to chronic pain. Identification of primate sensory neuron types is critical for understanding the cellular origin and heritability of chronic pain. However, molecular insights into the primate sensory neurons are missing. Here we classify non-human primate dorsal root ganglion sensory neurons based on their transcriptome and map human pain heritability to neuronal types. First, we identified cell correlates between two major datasets for mouse sensory neuron types. Machine learning exposes an overall cross-species conservation of somatosensory neurons between primate and mouse, although with differences at individual gene level, highlighting the importance of primate data for clinical translation. We map genomic loci associated with chronic pain in human onto primate sensory neuron types to identify the cellular origin of human chronic pain. Genome-wide associations for chronic pain converge on two different neuronal types distributed between pain disorders that display different genetic susceptibilities, suggesting both unique and shared mechanisms between different pain conditions.

## Introduction

The dorsal root ganglion (DRG) consists of a variety of neuron types, each tuned to detect and transduce different physical stimuli. These neuron types can broadly be divided into low-threshold mechanosensitive neurons responsible for sensing touch and high-threshold nociceptors, which are involved in pain, temperature and itch^1–4^. However, a comprehensive classification of DRG neurons is critical for understanding exactly how somatosensation works and for providing insights into the cellular basis for acute and chronic pain. Rodents represent the main species for studies on the cellular and molecular basis of nociception and the greatest insights with respect to molecular classification of neuronal types have been obtained from mouse, where single-cell RNA-sequencing (scRNA-seq) has led to a molecular taxonomy of existing types of sensory neurons^5–9^.

This has enabled the identification of molecular types representing richly myelinated A-fiber low threshold mechanoreceptors (LTMRs) and limb proprioceptors. The remaining neuronal types in the scRNA-seq are assigned as weakly myelinated or unmyelinated neurons. One of these is a C-fiber LTMR (C-LTMR) neuron type that expresses Vglut3 (*Slc17a8*) and tyrosine hydroxylase (Th) that likely is not involved in pain sensation^1–5^. Nociception is largely conferred through unmyelinated peptidergic C-fiber neuron types and a few lightly myelinated Aδ-nociceptors, a Trpm8 expressing cluster of neurons, as well as cell types marked by expression of Mrgprd, Mrgpra3 or Sst (named NP1, NP2 and NP3 types of neurons, respectively^8^). This molecular classification agrees remarkably well with previous studies based on myelination and conduction velocity, neurochemical features and termination patterns peripherally in the skin and centrally in the spinal cord and is also consistent with the known ontogeny of DRG neuron types^5^. As a result, there have been significant advances in understanding the cellular and molecular characteristics of sensory neurons found in mouse DRG.

Much less is known about characteristics of human DRG. Apart from information on size of the ganglia along the rostro-caudal axis, micro-anatomy including neuron size^10–12^ and electrophysiological characteristics^13–16^, the molecular characterization of human DRG is still limited to bulk RNA-sequencing^17–19^ and neurochemical analyses of gene products in a handful of studies^20^. Hence, the concordance of markers used in different studies and their relation to actual neuron types remain largely unknown. Nevertheless, by examining individual gene products, these studies suggest important species differences between human and mouse where for example Nav1.8, Nav1.9, P2×3 receptor and TRPV1 are present in both small and large neurons in human, but only small neurons in mouse, suggesting fundamental differences in molecular characteristics and principles of initiation and transduction of somatosensory stimuli between human and rodent^20^.

In humans, rare and drastic mutations that explain different types of congenital insensitivity to pain and erythromelalgia have been identified, such as for example *SCN9A* (Nav1.7), *NTRK1* (TRKA) and *SCN11A* (Nav1.9)^21–25^. In addition to these rare causing mutations, it is known that the genetic risk for chronic pain is due to common variations with small effect size^26^. Close to half of the risk of developing chronic pain are attributable to genetic factors^27–29^, including musculoskeletal pain conditions^28^. For musculoskeletal pain there is statistical evidence for a diverse set of genes involved, with a marked overrepresentation of genes expressed in neurons and functionally associated to neurotransmission, indicating a strong heritable component caused by altered functions of neurons^26^. Pleiotropy of single nucleotide polymorphisms (SNPs) among painful and non-painful conditions has also been shown^30^, even in human DRG^31^. It has recently become possible to connect genomic results to transcriptomics at the cellular level which allows for insights into the cell types which are fundamental for disorders. Thus, taking advantage of scRNA sequencing for mapping susceptibility genes to cell types, new insights have been made into the cell types involved for example in schizophrenia^32,33^, neuroticism^34^, intelligence^35,36^ and Alzheimer’s disease^37^, but such analyses have not been attempted for chronic pain conditions.

Knowledge on the molecular and cellular characteristics of primate DRG and their mouse correlates has critical implications for translating data from rodent models to human pain disorders^38,39^ and allows mapping genomic loci implicated in chronic pain onto specific primate somatosensory neuron types. Here we explore the cellular basis of somatosensation in the non-human primate and identify sensory cell types linked to human chronic pain.

## Results

### Molecular diversity of sensory neuron types in a non-human primate

We prepared DRG cell suspensions for scRNA-seq from adult Rhesus macaques using two different platforms (Fig 1a). First, cells from three macaques (two females and one male) were captured and sequenced using STRT-2i-seq. A total of 4,742 cells were sequenced and the reads were aligned to the macaque genome Mmu10 with gene names and annotations transferred from human (see methods). The data were merged and then clustered using the anchoring-based integration and graph-based clustering approach implemented in Seurat^40^. Using iterative rounds of clustering and quality control, we identified and removed non-neuronal cells, injured neurons and ambiguous cells, and finally merged clusters with highly similar transcriptomic profiles (Extended Data Fig.1a-g, see Materials and methods). The remaining 2,518 neurons formed nine separate clusters (Fig. 1b, Extended Data Fig.1g). For validation, this cleaned dataset was also analyzed using Conos^41^, an approach which Identifies multiple plausible inter-sample mappings and builds a joint graph of the datasets. This approach produced close to identical clusters to the original nine formed with Seurat (Extended Data Fig.1h, i; see Materials and methods). Gene expression patterns between the different animals showed near perfect positive correlation indicating high similarity of inter-individual transcriptome profiles (Extended Data Fig.1j). The analyzed neurons contained 5,687 genes and 38,624 unique transcripts per cell on average, expressed neuron and sensory neuron specific genes (*RBFOX3, SLC17A6*) throughout with limited expression of satellite-glia genes (*FABP7, APOE*) and showed unique gene expression profiles (Extended Data Fig.1k, l; Fig. 1c, d).

**Fig. 1.**
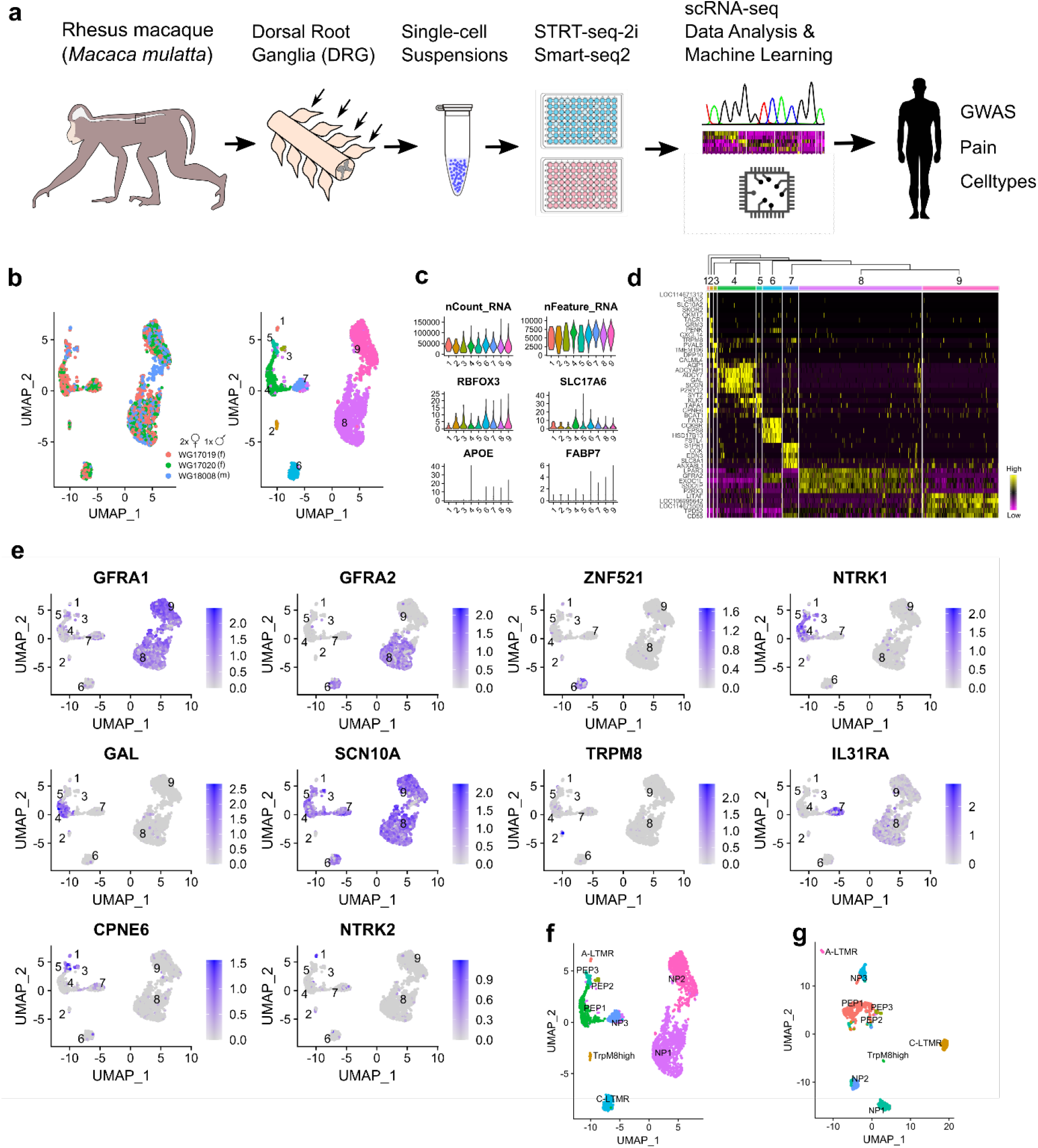
Somatosensory neuron clusters in the macaque DRG. (a) A schematic view of the workflow. (b) UMAP plots showing (left) the contribution of individual animals to STRT-2i-seq clusters and (right) final cluster numbers (m=male, f=female). (c) Violin plots showing total counts of unique transcripts, detected genes, and transcript counts for neuronal and satellite glia marker genes in the neuronal clusters. Y-axes shows detected genes per cell for nFeature_RNA, all others are raw UMI counts. (d) A hierarchically organized heatmap with the five most specific genes (by p-adj) for each cluster. (e) UMAPs showing mouse canonical marker gene expression in the STRT-2i-seq macaque clusters. (f) STRT-2i-seq macaque clusters named after most likely mouse counterparts. (g) Named Smart-seq2 clusters after label transfer from the STRT-2i-seq data.

After clustering, we used canonical mouse DRG neuron markers to assign tentative identities for the clusters based on their likely mouse counterparts (Fig. 1e, f): NP1 (cluster 8) and NP2 (cluster 9) were named based on the combination of *GFRA1* and *GFRA2* expression; C-LTMRs (6) were assigned using *GFRA2* and *ZNF521* (*Zpf521* in the mouse); *NTRK1* and *GAL* were used to identify PEP1 (4); *SCN10A* and *TRPM8* suggested the identity of the TrpM8high (2) cluster (negative for *SCN10A*); *IL31RA* expression was used for naming NP3 (7); *CPNE6* together with *NTRK2* was used to assign putative A-LTMRs (1), and *CPNE6* together with *NTRK1* and *SCN10A* were used to assign the PEP2 (3) cluster. The final cluster (5) also expressed *CPNE6, NTRK1* and *SCN10A*, and was named PEP3.

A second dataset was prepared from 5 female macaques using Smart-seq2 technology. After clustering and cleaning steps (Extended Data Fig.2a-d) these data included 1038 neurons showing >480,000 counts and >13,500 detected genes per cell on average (Extended Data Fig.2e, f). Tentative neuron identities for this dataset was assigned in a supervised manner using the STRT-2i-seq data as a reference (Fig. 1f, g; Extended Data Fig.2g). Interrogation of marker gene expression used for tentative cell type assignment showed identical patterns between the two macaque datasets (Extended Data Fig.2h), thus representing an independent identification of the cell types identified in the STRT-2i-seq data. For full marker lists from both scRNA-seq datasets, see Extended Data Tables 1 and 2. An interactive web resource for browsing the datasets is available at: https://ernforsgroup.shinyapps.io/macaquedrg/.

### A consensus on mouse DRG neuron types across datasets

We wanted to further leverage the vast knowledgebase of mouse DRG neuron types in our investigation of primate DRG neurons. For this, we used two major scRNA-seq datasets from previously published mouse studies, referred hereafter as the Zeisel and Sharma datasets^7,9^. These studies identify similar number of cell types, but it is not known if the same kinds of neurons were identified and furthermore, the studies use different nomenclature. We therefore first identified the corresponding cell types between the datasets. Using the label transfer method implemented in Seurat, we transferred labels from Sharma over to the Zeisel data (Fig. 2a, b; Extended Data Fig. 3a) and then also named the Zeisel types using Usoskin nomenclature^8^ (Fig. 2c). We then repeated the label transfer from Zeisel to Sharma data and named the Sharma types also using Usoskin nomenclature (Fig. 2 d-f, Extended Data Fig. 3b). Finally, we repeated the label transfer from Zeisel to Sharma both ways using Usoskin labels for the Zeisel data, and observed identical results (Extended Data Fig. 3c, d).

**Fig. 2.**
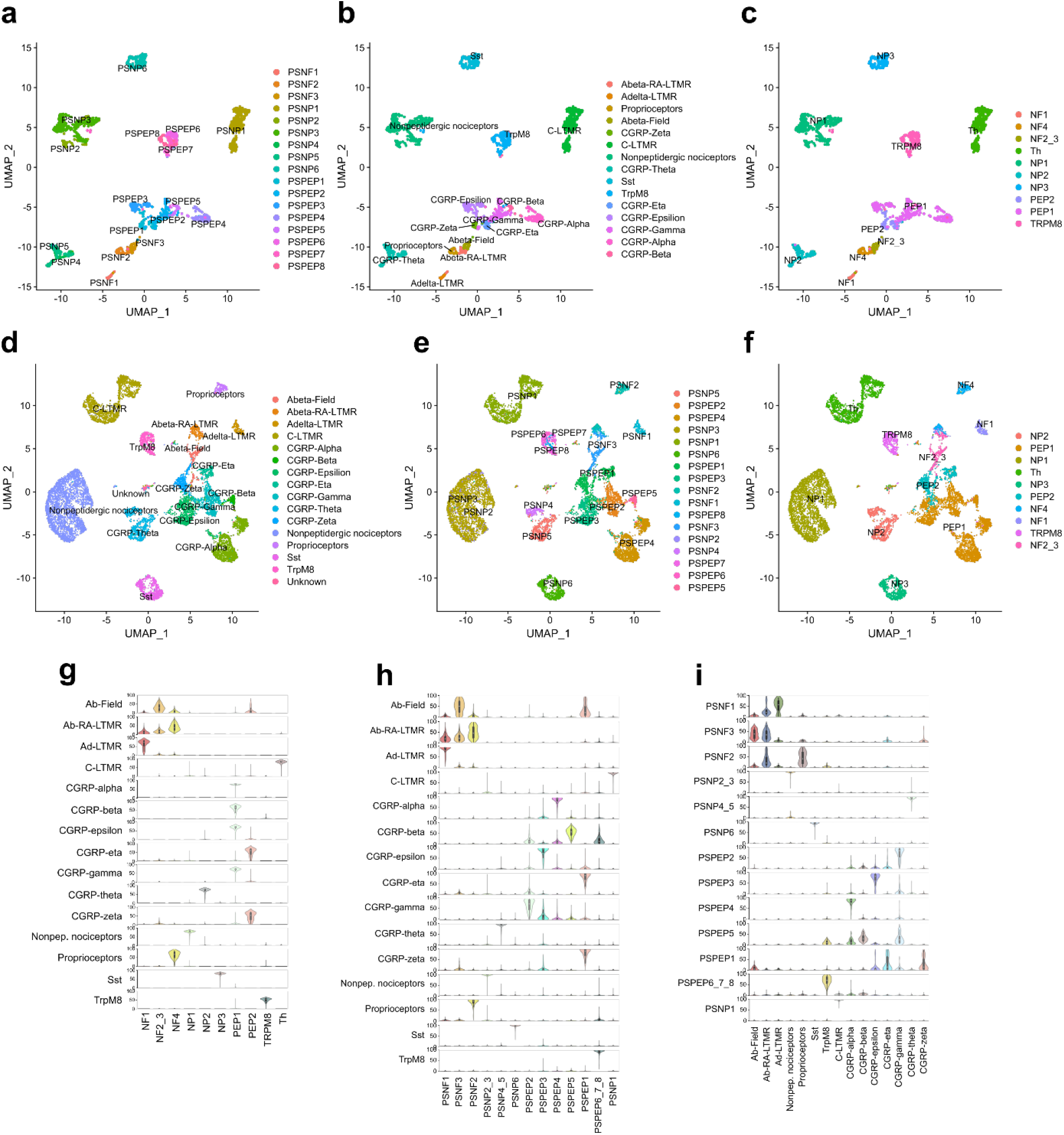
Consensus of mouse DRG neuron types across datasets and nomenclatures. (a-f) UMAPs showing (a) Zeisel et al. types with Zeisel nomenclature, (b) Zeisel et al. data after label transfer from Sharma et al. data, (c) Zeisel et al. data with Usoskin nomenclature, (d) Sharma et al. data with original nomenclature (e) Sharma et al. data after label transfer from Zeisel et al. data and (f) Sharma et al. data with Usoskin nomenclature. (g) Probability scores of Sharma et al. types against Zeisel trained module (performed with Usoskin nomenclature). (h) Probability scores of Sharma et al. types against Zeisel trained module (performed with Zeisel nomenclature). (i) Probability scores of Zeisel et al. types against Sharma trained module (performed with original nomenclature from Sharma et al 2020).

As an independent method to establish the similarity between cell types identified in the different studies, we employed neural-network based probabilistic scoring modules for learning cell-type features between datasets. We trained the modules using both mouse datasets and using all three nomenclature versions (Zeisel and Usoskin nomenclatures for Zeisel data; Sharma nomenclature for Sharma data) (Extended Data Fig. 4a-i). We then tested the performance of these modules in finding the corresponding cell types between the datasets and found that the modules detect with high probability cell types across the different datasets and that the cell-type assignments concur with the results obtained by label transfer. For the rest of our analyses we used the Usoskin type nomenclature as the default naming. Taken together, the results confirmed a one-to-one relationship between nociceptor types identified in^7,9^ although the Aδ-nociceptors of Zeisel, were split into two subtypes in Sharma (Fig. 2a-i). Because Usoskin nomenclature performed with the least noise and greatest prediction scores, we used this nomenclature for the rest of our analyses.

### Overall cross-species conserved strategy for somatosensation

We proceeded to use the probabilistic neural-network machine learning approach to evaluate whether the neuronal basis of somatosensation is conserved between macaque and mouse, and to validate our tentative assignment of cross-species neuron correlations. For this purpose, we generated the probability score for each macaque cluster to the mouse neuron types using both the macaque STRT-2i-seq and Smart-seq2 datasets. Each of the macaque clusters showed similarity to the previously assigned mouse neuron types (Fig. 3a, b, Extended Data Fig. 5). These results also indicated that the macaque A-LTMR cluster consists of cells corresponding to the lightly myelinated Aδ-LTMR type. Our original annotation of corresponding macaque-mouse neuron types was consistent with the expression of *PRDM12* in all nociceptors and the mechanosensory channel *PIEZO2* in A-LTMRs, C-LTMRs and NP1 (Fig. 3c). Interestingly, the mouse proprioceptor marker *PVALB* was expressed in the macaque PEP2 neurons. PEP3, the other macaque neuron type showing similarity to mouse PEP2, expressed lower levels of *TRPM8*, moderate *PIEZO2* and differed from the macaque PEP2 by expressing *KIT, SCGN* and lower levels of the heat sensitive channels *TRPV1* and *TRPA1* (Fig. 3d). When compared to the mouse PEP2 subtypes in Sharma et al., PEP3 showed higher probability to CGRP-eta over GCRP-zeta and PEP2 to GCRP-zeta over CGRP-eta (Extended Data Fig. 5a, b), indicating two types of Aδ-nociceptors.

**Fig. 3.**
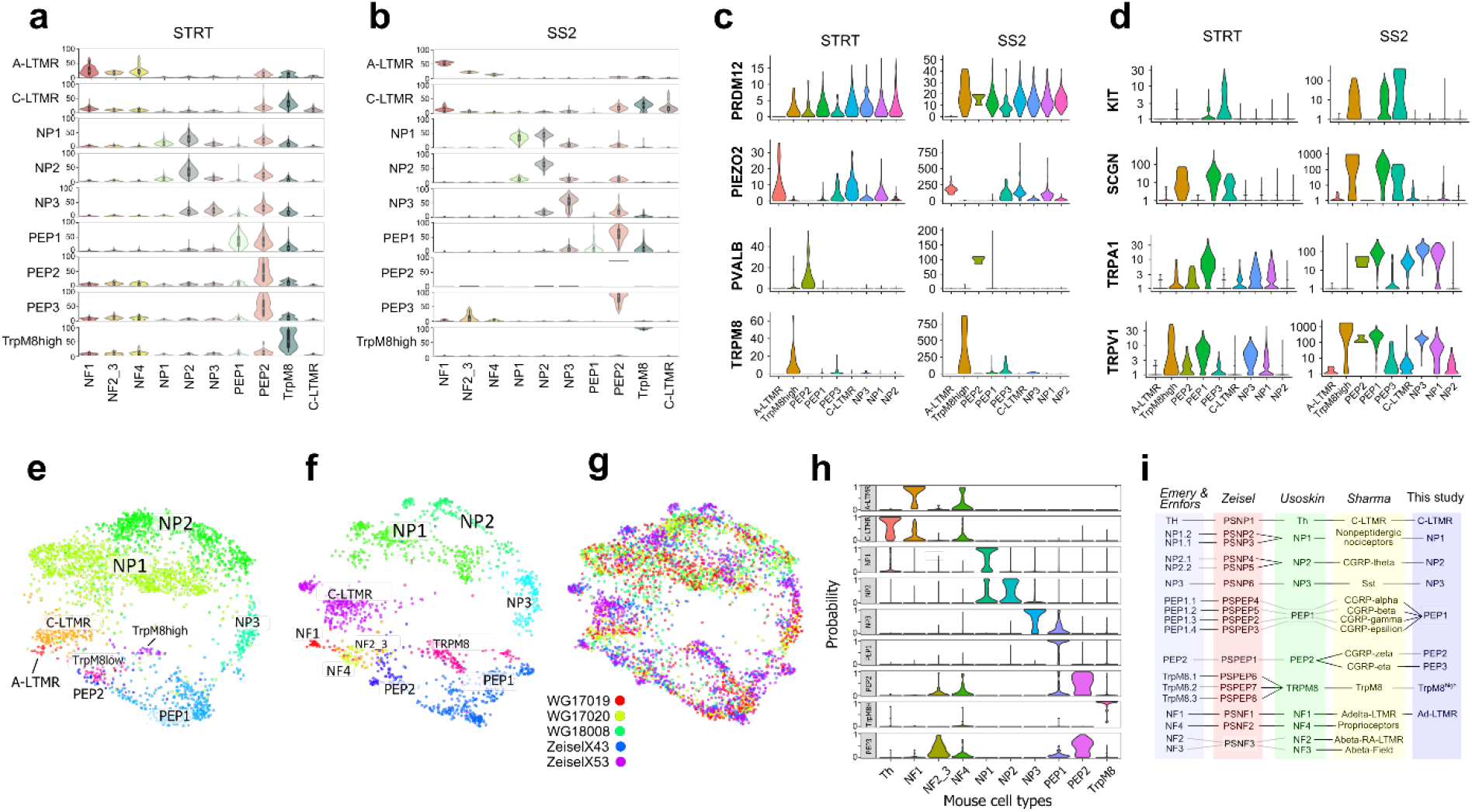
Identification of primate correlates of mouse neuron types. (a, b) Violin plots showing prediction percent probability scores for each macaque cluster against established mouse DRG cell types trained with Usoskin nomenclature. (a) STRT-2i-seq, (b) Smart-seq2. (c) Violin plots of markers expressed in macaque clusters. Y-axis, raw UMI counts for STRT-2i-seq and counts for Smart-seq2. (d) Violin plots showing genes differentially expressed between macaque PEP2 and PEP3. (e-g) LargeViz plot showing cross-species clustering of macaque STRT-2i-seq and mouse Zeisel et al. datasets on a shared plot. (e) macaque, (d) mouse, (g) merged plot (legend shows origin of samples, WG = macaque, Zeisel = mouse). (h) Probability plot for each macaque neuron type against mouse neuron types. (i) Synopsis of the corresponding DRG neuron types between mouse and macaque datasets and nomenclatures^5,7–9^.

To gain further confidence in our cross-species analyses we performed co-integration of our macaque STRT-2i-seq data with mouse data from Zeisel et al. ^9^ using Conos. Here, the previously assigned macaque clusters showed close positioning to homologous mouse clusters on a joined cross-species clustering graph (Fig. 3e-g). This strong cross-species association was also apparent on the probability profiles after label propagation from mouse clusters to individual macaque neurons.

All macaque clusters showed close to one-to-one correspondence to individual mouse clusters with PEP2 and PEP3 having the strongest association to the mouse PEP2 cluster (Fig. 3h). Combined, these results evidence a strong cross-species association of sensory neuron types, indicating that the overall cellular basis for somatosensation is conserved between mouse and macaque (Fig. 3i).

The macaque neuronal types were validated *in vivo* by triple *in situ* hybridization (Fig. 4a, Extended Data Fig. 6). *SCN10A* was used as a general marker for nociceptors together with cluster-specific markers or a combination of markers defining only one cluster. For some clusters, negative markers were used to rule out types other than the one under analysis (see Materials and Methods).

**Fig. 4.**
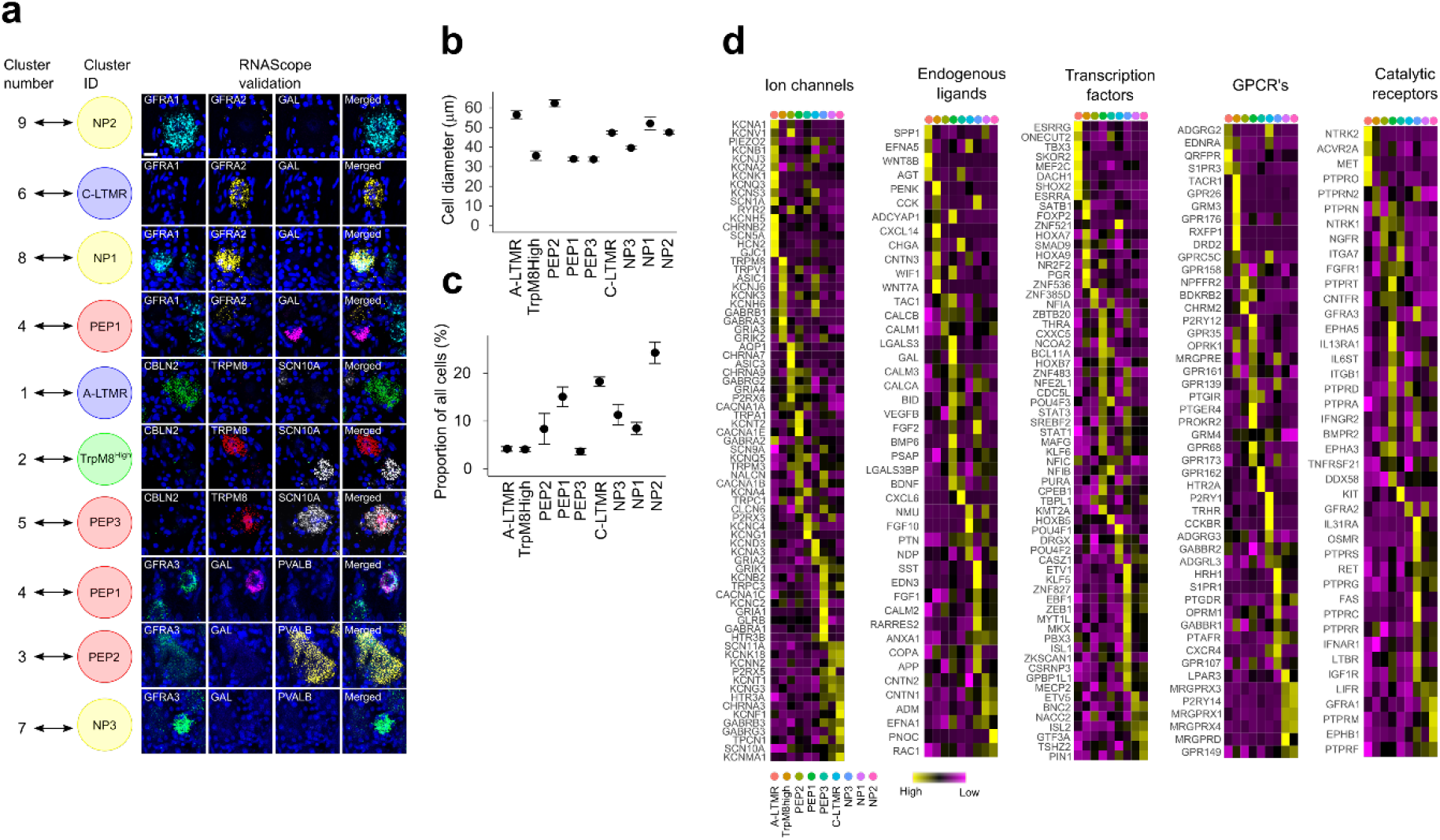
In-vivo validation and gene family expression in the macaque DRG neuron types. (a) Naming the macaque clusters after the closest corresponding mouse cell types (left) with RNAScope validations (right). Scale bar = 20µm. (b) Diameter of neuronal type populations in lumbar DRGs, mean +/- SEM, n = 3 animals. (c) Proportion of each neuronal type among all neurons based on in vivo data (mean +/- SEM, n=3 animals) in lumbar DRGs. (d) Heatmaps of gene expression profiles for selected gene families in the macaque STRT-2i-seq DRG neuron clusters.

In addition, we analyzed the soma size distribution and percentage contribution of each cell type in the DRG from the *in vivo* data (Fig. 4b, c). Finally, we interrogated each of the neuron-types for unique expression patterns for transcription factors, ion channels, G-protein couple receptors (GPCRs), catalytic receptors and endogenous ligands, including neuropeptides (Fig. 4d). This revealed for example the expression of multiple GPCR’s related to exogenous defense and cholestatic itch in NP1 and NP2 (*MRGPRX1-4*), and to histaminergic itch and inflammatory lipids in NP3 (*HRH1, S1PR1*) and NP1 (*LPAR3*).

### Species differences and similarities in gene expression

Our above results confirm an overall existence of neuronal correlates between the mouse and a primate, nevertheless, important divergences could still exist when examining expression of individual genes within each of the different mouse-macaque correlates. In order to provide a more comprehensive map of the molecular conserved and divergent features of somatic sensation and pain between the mouse and the macaque, we compared gene expression patterns between the species (Fig. 5a-c and Extended Data Table 2). Because strategies to identify new molecular targets for development of analgesic drugs often are focused on genes expressed uniquely in the neuron type(s) causative of pain, we examined the presence of genes within the different neuron types to identify conserved transcriptional programs between species as well as sets of genes that are expressed in highly species-specific manner between corresponding cell types. In such analyses, false negatives can confound the results. We therefore first examined the reliability of the individual STRT-2i-seq and Smart-seq2 datasets in side-by-side analyses of cell type specific expression patterns of mouse-macaque shared genes, macaque-specific genes and mouse specific genes, which showed high reproducibility across different platforms (Extended Data Fig. 7a-c). We thereafter combined the datasets for mouse^7,9^ and macaque (STRT-2i-seq and Smart-seq2) to obtain integrated datasets for mouse and macaque. Analysis of this dataset revealed the existence of robust conserved molecular features between mouse and non-human primate (Fig. 5a). For example, neuronal type specific mouse-macaque conserved features of NP3 neurons include *SST, JAK1, IL31RA, OSMR* and *S1PR1* (N = 36 genes) and C-LTMRs *P2RY1, EXOC1L, KCND3, IQSEC2, OSBPL1A* and *FXYD6* (N = 60 genes). The largest cell-type specific shared gene program was found between mouse and macaque Aδ-LTMRs (N = 126 genes) whereas NP1 and NP2 both had conserved features of less than 30 genes (N = 26 and 27, respectively). However, we also identified cell type specific gene expression that were species specific and these existed both in mouse and macaque (Fig. 5b, c; Extended Data Table 3). As examples, species differences included in NP3 the specific expression of *TDRD1, EDN3* and *GRIA1* in macaque and *NPPB, HTR1F* and *NTS* in mouse. For C-LTMRs, *TH* and *RARRES1* are specific for mouse, whereas *HSD17B13* and *CCKBR* are specific for macaque. These species-specific expression patterns need to be considered when translating results obtained in rodents to primates.

**Fig. 5.**
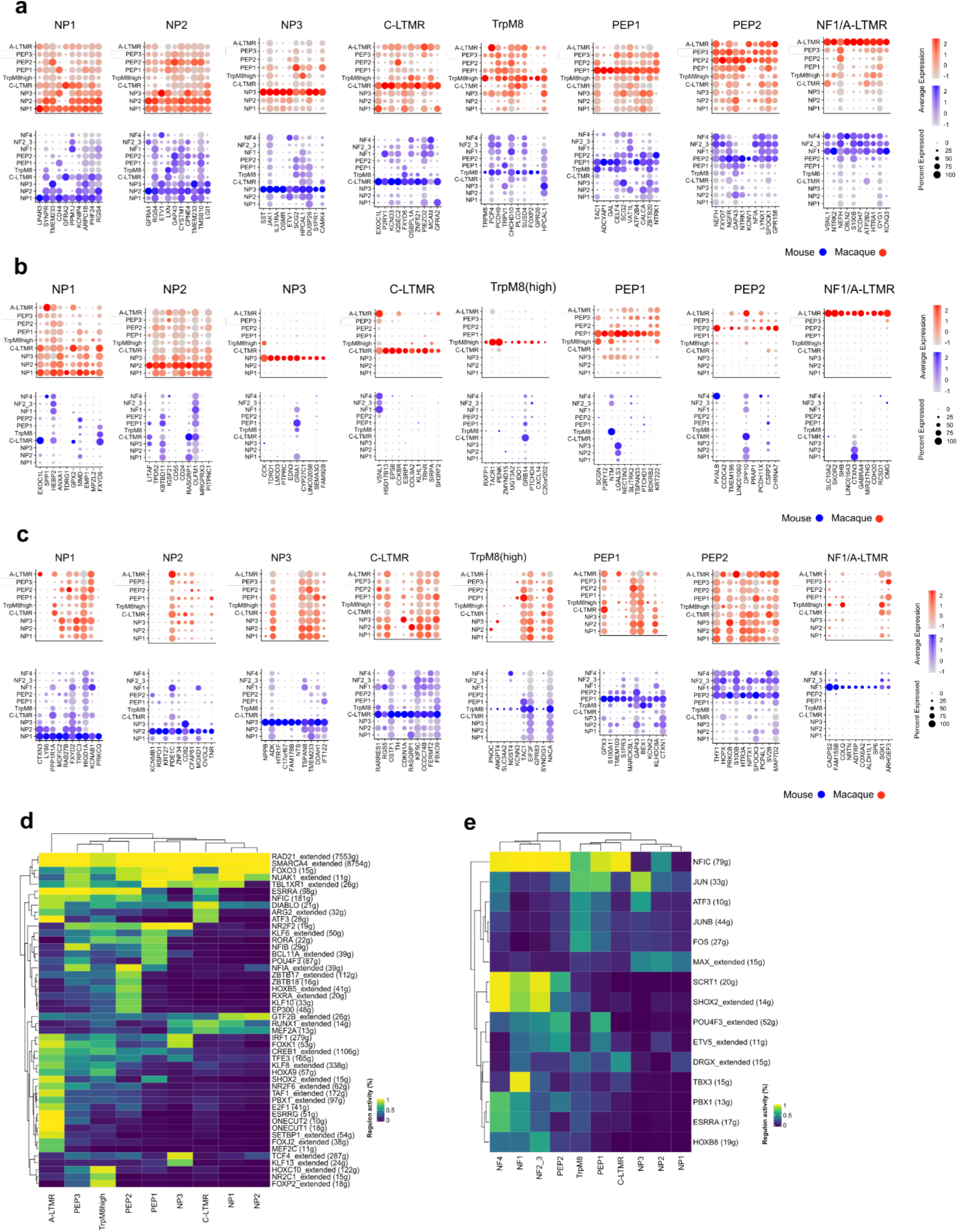
Comparison of gene expression profiles across species. (a) Dot plots showing ten most specific shared markers between closest related cell types between the species. (b) Dot plots showing ten most macaque specific markers for each corresponding cell type pair. (c) Dot plots showing ten most mouse specific markers for each corresponding cell type pair. Dot sizes in (a-c) correspond to percentage of cells expressing the gene in the cluster and color scale indicates log2FC. (d) Heatmap showing binary regulon activity in each of the macaque DRG cell types. (e) Heatmap showing binary regulon activity in the mouse DRG cell types. In (d, e) the number of target genes is indicated in the parenthesis.

We further performed supervised computational screens to find gene families whose differential expression could reliably distinguish similar cell types within and across species^42^. A set of over 1,500 gene families from HGNC (HUGO Gene Nomenclature Committee) was used for these screens (Extended Data Table 4). Within each species the top performing families included ion channels, G-protein coupled receptor families, cell adhesion molecules and others (Extended Data Fig. 8a, Extended Data Tables 5 and 6). Across species the top performing families included voltage-gated ion channels, G-protein-coupled receptors and neuropeptides (Extended Data Fig. 8b, Extended Data Table 7), suggesting that sensory neuron identities culminate mostly on genes of these families even across species. On cell type level (Extended Data Fig. 8c, Extended Data Table 7), A-LTMR/NF1 neurons were identified nearly perfectly by many gene families, for example by ion channels and cell adhesion molecules (CAMs). For the peptidergic C-fibers (PEP1), neuropeptides/receptor ligands were the highest performing family, as expected. On the other hand, NP1 and NP2 scored poorly in comparison to all other types, suggesting that these neuron types have diverged the most in their gene expression signatures between mouse and macaque. As a final comparison between the species, we used SCENIC^43^ to identify gene regulatory networks formed from master transcription factors and their gene targets (i.e. regulons) in all DRG cell types of both macaque and mouse (Fig 5d-e). We found for example the known fate determining regulons driven by *SHOX2, RUNX1* and *FOXP2* in mouse^9,44–46^ and predicted several new species unique as well as cross-species conserved regulons determining cell-type identities (see Extended Data Tables 8 and 9 for genes).

### Genetic association of neuronal types contributing to human pain states

The identification of the cellular and molecular basis for somatosensation and pain in non-human primate enables to determine the contribution of different primate neuron types in human chronic pain states. We therefore used human genetic data to explore how each cell type in the macaque DRG connects to painful phenotypes in humans by employing genome-wide association studies (GWAS). To do so, we used a large cohort made available by the UK Biobank project^47,48^, where we assessed chronic pain from self-report at eight body sites (Fig. 6a). Among the nine neuron types identified in the macaque DRG, we found that common variants associated with chronic pain sites mapped to sets of genes that were specifically expressed in two neuron types in the STRT-2i-seq dataset. We found enrichment of headaches, facial, neck and shoulder, stomach, and hip chronic pains partitioned heritability in PEP1 neurons (*P*_FDR_=16%, each), while NP2 neurons were associated most significantly with the heritability of chronic back pain and hip pain (*P*_FDR_=5%, 8%, respectively) (Fig. 6b, Extended Data Table 6). Thus, heritability of hip pain was significantly enriched in both PEP1 and NP2 neuron types while heritability of all other pain sites was significantly enriched in only one neuron type of the macaque DRG. Since the epidemiological prevalence of chronic pain patients to report more than one body site is high, the signal attributed to, for example, subjects with hip pain may also have back pain, so it is not clear where the association signal is deriving from. To control for this co-morbidities, we performed new full GWASes for each of the pain site (row in Extended Data Fig. 9b, c), then removed one by one all comorbid pain sites (column in Extended Data Fig. 9b, c) in all GWASes for PEP1 and NP2 neurons. Although some statistical power is lost in this analysis due to reduced size of chronic case participants, partitioned heritability in PEP1 was confirmed in most GWASes (facial, neck/shoulder, stomach/abdomen and hip pain), while association with back pain remained negative (row in Extended Fig. 9b). However, significance of PEP1 for headaches was lost for all GWASes. Furthermore, back pain and hip pain remained significant for NP2 neurons in all GWASes when excluding other pain sites (row in Extended Fig. 9c). Our results show that seven of the nine neuron types were not associated with any chronic pain sites, and hence the two neuron types, PEP1 and NP2, together represent the main enrichment of musculoskeletal pain heritability. A meta-analysis across all pain sites for each cell type (Fig. 6c) consistently found that both PEP1 (*P*=5.4⨯10^−3^) and NP2 (*P*=3.9⨯10^−3^) displayed significant enrichment across all pain sites.

**Fig. 6.**
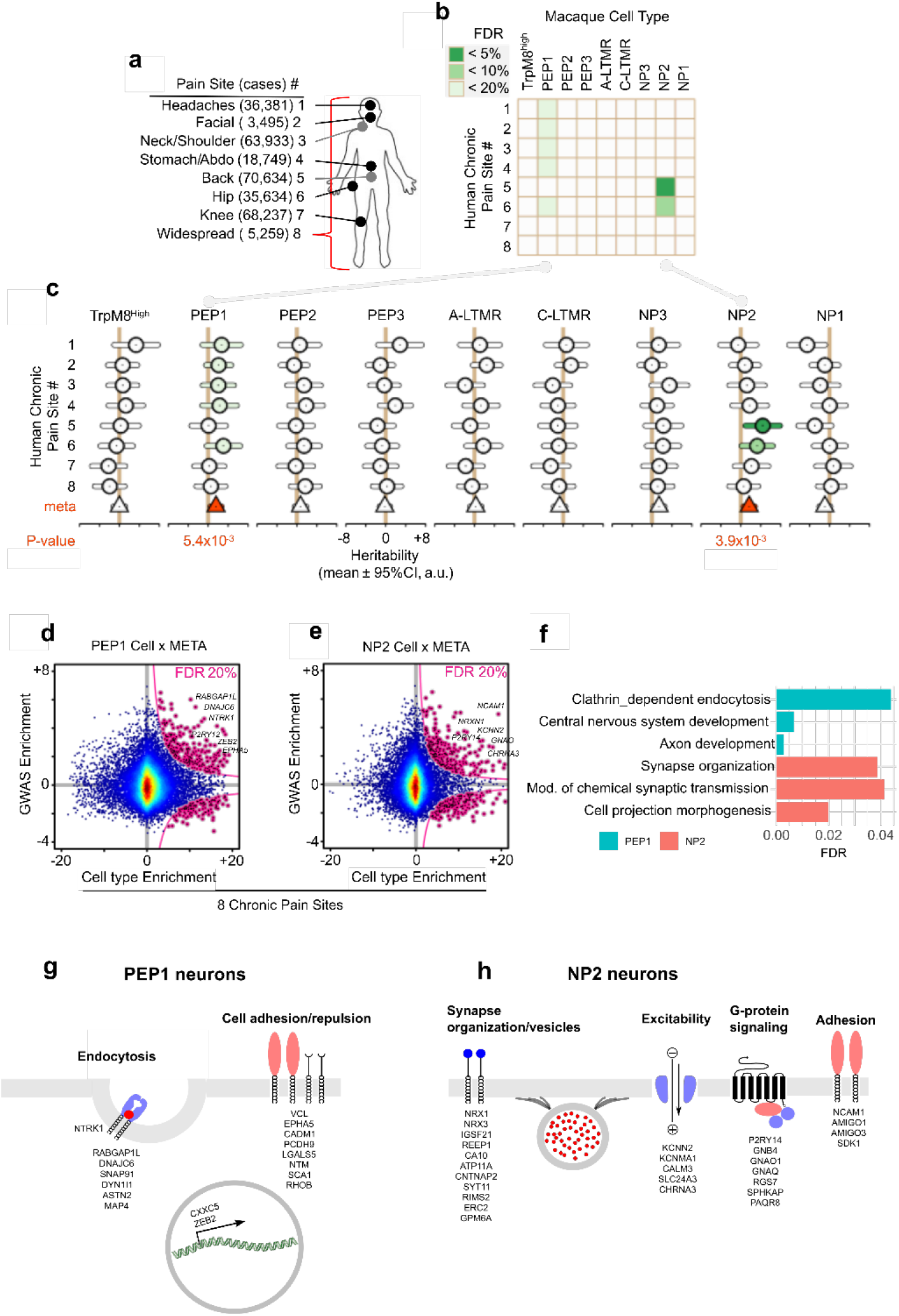
Contribution of macaque DRG cell types to partitioned heritability of human chronic pain sites. (a) UK Biobank chronic pain sites mapped to the human body. Number of chronic case participants shown in parentheses. (b) Heatmap of FDR-corrected p-values for enrichment in partitioned heritability of each macaque DRG cell type to each human chronic pain site. (c) Forest plot of partitioned heritability estimates for each macaque DRG cell type contributing to chronic pain. Shown are heritability coefficient estimates (circles) and their 95% confidence intervals (bars) for each pain site. Fixed-effect, standard error weighted, meta-analyzed heritability estimates (triangles) also shown, colored red when significant at Bonferroni-corrected level (corrected for nine cell types). (d,e) Top genes in type-specific cells contributing to specific chronic pain sites. Top genes highlighted in a scatter plot (pink, left). Scatter plots show human GWAS enrichment of gene (Y axis) as a function of macaque cluster enrichment (X axis). (f) Bar plot of top pathways for PEP1 and NP2 cell types in the meta-analysis of the 8 chronic pain sites. (g, h) Schematic illustrations of pathway-related genes in (g) PEP1 and (h) NP2 neuron types.

We next tested if these results were reproducible in the independent scRNA-sequencing dataset obtained by the Smart-seq2 protocol. Consistent with the STRT-2i-seq dataset, we found enrichment of stomach, hip, and neck and shoulder chronic pain partitioned heritability in PEP1 neurons (*P*_FDR_=10%, 10%, 13%, respectively), while NP2 neurons were associated most significantly with the heritability of chronic back pain, hip pain and knee pain (*P*_FDR_=10%, each) (Extended Fig. 9d). Thus, heritability to pain was consistently assigned to the same neuronal types using the STRT-2i-seq and Smart-seq2 datasets, although significance of PEP1 to headaches and facial pain was lost in the latter.

In order to identify functional pathways and genes whereby human heritability contributes to chronic pain in the different neuronal types, genes were ranked by exclusivity of neuronal cell type expression to establish enrichment scores. Human GWAS enrichment scores were mapped to the macaque single cell expression enrichment to identify top genes in type-specific cells contributing to chronic pain sites in PEP1 and NP2 neurons, revealing hundreds of genes (Fig. 6d, e, Extended Data Table 10). These were thereafter used to identify cellular pathways that confer vulnerability to chronic pain (Fig. 6f). Combined, these results show that heritability of chronic pain at different sites is associated with each of the two major pain neuron types through different biological pathways (see Extended Data Table 10). Pathways contributing to chronic pain in PEP1 neurons included “clathrin-dependent endocytosis”, “central nervous system development” and “axon development” with an enrichment of proteins involved in the process of endocytosis, cell adhesion as well as a few transcription factors (Fig. 6f, g). In contrast, pathways in NP2 neurons included “synapse organization”, “chemical synaptic transmission” and “cell projection morphogenesis” and were dominated by genes associated with organization of the synaptic membrane and its vesicles, ion channels participating in excitability, G-protein signaling and cell adhesion (Fig. 6f, h). Thus, the two neuron types contribute to chronic pain via distinct pathways.

## Discussion

The human DRG like the mouse contain neurons with different histochemical and electrophysiological features^13,14,14–16,49,50^. The identification of the molecular types of primate somatosensory neurons addresses the longstanding question whether cell types involved in somatosensation is conserved between rodents and primates. We conclude that the mouse^7–9^ and the Rhesus macaque largely share molecular neuron types which using mouse genetics have been functionally identified as A-LTMR involved in touch and proprioceptive sensation^1^, C-LTMRs involved in the affective aspect of pleasant touch^5^, C-cold thermoreceptors (TrpM8^high^), Aδ fast mechanical nociceptors involved in pinprick pain (PEP2)^51–54^ and mechano-heat C-nociceptors (PEP1), as well as “non-peptidergic” neuronal types (NP1, NP2, NP3) known in mouse to be involved sensing noxious mechanical threshold and itch sensation^55–57^. Although the neural network predictions score was low, macaque PEP3 appears to be an Aδ−fiber fast nociceptor which correspond to mouse CGRP-eta type in Sharma nomenclature, which is a subtype of the mouse PEP2 type in the Usoskin nomenclature.

Overt differences in the overall cellular basis for nociception between mouse and macaque largely relates to NP1 and NP2 neurons. In the mouse, NP1 is involved in detecting pricking mechanical stimuli and β-alanine induced itch through the Mrgprd receptor, but not thermal sensation. In the macaque, the expression of the heat-activated channel TRPV1 in NP1 neurons, in addition to TRPA1, suggests a broader function than in the mouse. The mouse NP2 neurons express histamine and chloroquine receptors and ablation of these neurons (Mrgpra3^+^ neurons) specifically affects histamine-dependent and histamine-independent itch, but not acute noxious heat, cold, or mechanical pain^56^. NP2 neurons have therefore been considered to be dedicated itch neurons in rodents^58^. However, this neuronal type was recently found to code for both itch and pain, with itch behavior induced by metabotropic Gq-linked stimulation and pain behavior through fast ionotropic stimulation^59^, suggesting that the same neuronal populations can drive distinct sensations in a stimulus-dependent manner. In addition, based on the molecular profiles, macaque NP2 neurons appear partly different from mouse as histamine receptor HRH1 is low in macaque NP2 neurons. Other known functional stimuli detectors found in these cells are MRGPRX1-4, which are also expressed in NP1 neurons. MRGPRX1-4 are promiscuous low-affinity receptors involved in non-histaminergic itch, for example chloroquine and pruritogenic peptides^58^. Primate NP1 and NP2 neurons may thus have at least partly different functions than in the mouse.

Even though the exact neuronal basis for human chronic pain is unknown, insights have been obtained through the identification of genes causing congenital insensitivity to pain^22,60^. While most of the genes causing painless phenotype are abundantly expressed in all DRG neuron types, some display restricted expression patterns, thus opening for linking neuronal types to phenotype. These includes congenital insensitivity to pain by mutations in *SCN9A* (Nav1.7), *SCN11A* (Nav1.9) and *NTRK1* (TRKA)^21–25,61^ *and PRDM12*^62^. In contrast to mouse which display an enriched expression in nociceptors, macaque *SCN9A* is broadly expressed at similar levels in all neuronal types, while *SCN11A* expression is more similar to mouse with expression at varying levels in all unmyelinated neuronal types (C-LTMRs, PEP1, NP1-3,) with very low levels in TRPM8^high^, myelinated nociceptors and A-LTMRs (see https://ernforsgroup.shinyapps.io/macaquedrg/ for interrogation of gene expression). *NTRK1* is largely confined to macaque TRPM8^high^, PEP1, PEP2 and PEP3 neuronal types, with lower levels in the other nociceptors. *PRDM12* expression is consistent with mouse, appearing in all macaque neurons except Aδ-LTMRs. Thus, although it is not possible to pinpoint the exact neuronal types, it seems based on expression of these causative genes for human monogenic pain insensitivity disorders that PEP1, PEP2, PEP3, TRPM8^high^ represent important neuronal types for nociception. However, it cannot be excluded that neuronal types sufficient for driving chronic pain might partly involve neuron types other than those required for nociception.

Previous GWAS studies have uncovered genome-wide significant genes that contributes to the heritable risk of chronic pain. Recent methods^32,33^ allow the integration of GWAS and scRNA-seq data to map cell types contributing to disease through the testing the enrichment for cell type-specific expression of genes with nearby risk SNPs. Such analyses consider the heritability carried by all common SNPs, linking them to nearby genes, rather than focusing only on genome-wide significant genes. Using this methodology, we linked GWAS results of several human chronic pain sites to specific neuronal types in the primates.

Significance for cell type specific contributions reported in previous studies^32,63^ showed stronger association than that reported in this work (FDR in the range of 5-20%). However, the estimation for the heritability of pain ranges from 2-10%^64–66^ with 7.6% for chronic back pain^67^. In a comparative study of heritability between different classes of diseases, it was shown that mental health disorders show higher heritability estimates than self-reported pain phenotypes^66^. Furthermore, all chronic pain GWAS in the UK Biobank displayed inflated genomic control parameters (λ_GC_ >> 1) while at the same time an LD regression score intercept close to 1, indicating that the inflation’s origin is highly polygenic in nature. Because of these two effects combined (lower heritability and higher polygenicity), a tissue- or cell-type specific contribution to chronic pain would be smaller in comparison with other diseases like schizophrenia. On the contrary, it is perhaps remarkable that some specific cell types show significance, particularly because the burden also distributes to cell types other than sensory neurons. However, the strength of our analysis is that it considers the accumulated contribution of all small effect sizes in sensory neurons distributed across the human genome. Thus, we believe that the significance observed is in the range of expectation. Furthermore, we show that the association to the identified neuronal types emerge from multiple pain GWAS and in addition was reproduced in an independent dataset using a different sequencing platform.

The results show that musculoskeletal pain genetics is well represented in the DRG. Our findings reveal a connection to two of nine types of neurons to multiple chronic pain sites, namely PEP1 and NP2. However, there is an overlap of individuals being included in some of these pain groups and if a cell type is enriched in multiple pain GWAS, there is a risk of reporting the false association that is driven by phenotypic comorbidity but not a true genetic association. Iterative removal of one pain site at a time in the GWAS and examining if the significance to the other pain site remains, indicate that most of the GWAS signals observed is attributed to the annotated pain sites themselves. These results point towards a common underlying pain vulnerability regardless of the body site where pain is manifested. PEP1 for headache could not be confirmed in this analysis. Thus, assignment of contribution of PEP1 cell type to headaches might be due to comorbidity of other pain sites with headaches. This suggest that musculoskeletal pain and headache are unique experiences caused by different neuronal mechanisms, which is in line with reports that in congenital insensitivity to pain phenotypes, there are painless cases with the only pain felt consisting in tension headaches^68^ and furthermore, is consistent with findings that there is a shared genetic factors in conditions manifesting chronic pain except for migraine^69^.

Apart from headache, the different pain sites suggest the involvement of one of two main neuron types PEP1 or NP2 in all cases but hip pain, which is associated with both. Thus, these results suggest that different neuron types are associated with different chronic pain sites. This opens for the question whether the different types of pain could be location dependent, thus depending on segmental levels of DRGs or trigeminal ganglion neurons in the facial region rather than neuronal type dependency. However, analysis of somatosensory neurons across the rostro-caudal axis of the mouse including DRG^7–9^, jugular ganglion^70^ and trigeminal ganglion^7^ reveals that the neuronal strategy of somatosensation is shared regardless of body location. Thus, similar types of somatosensory neurons exist in the DRG as in jugular and trigeminal ganglion. Because of this, we assume that rostro-caudal differences in pain sites should not affect the identification of involved cell-types. This motivated us to use macaque DRG to predict the heritable risk genes at different pain locations in the human, including trigeminal area pain. While mouse models have not specifically addressed the contribution of NP2 neurons to chronic pain, the contribution of PEP1 neurons to pain is well established. For example, ablation of CGRP^+^ neurons, which include the PEP1 neurons, results in the loss of noxious heat sensation as well as inflammatory and neuropathic heat hyperalgesia in the mouse^71^.

Largely different genes contribute to chronic pain in PEP1 and NP2 neurons. In previous human association studies for musculoskeletal chronic pain there is a marked enrichment of genes involved in neurotransmission but also for example immune function, metabolic processing, skeletal tissue differentiation and hormone signaling pathways^26^. In this study, we expected to capture the signal associated with the heritable risk in the different somatosensory neurons. The results reveal unique vulnerability pathways at play during pain chronification in the different neuronal types, suggesting that the causal mechanisms might be different between chronic pain conditions. While there is an enrichment of risk genes in PEP1 neurons that belongs to clathrin-dependent endocytosis and axon and nervous system development, NP2 neurons display an enrichment in synaptic organization and transmission and cell projection morphogenesis. This does not inform that these pathways are the only ones present in the different sensory neuron of a particular population. Instead it shows that expression of, for example, specific members of cell adhesion/repulsion genes carrying nearby variations with significant heritable risk to chronic pain are enriched in PEP1 neurons. Analysis of the underlying genes with enriched heritability reveal a common pattern related to a susceptibility of neuronal connectivity, although with different functional classes of genes in different neurons. Thus, we conclude that the results support the notion that the major genetic risks expressed in somatosensory neurons are carried in genes involved in structural and functional connections of the neurons and thus, impacts on neurotransmission. The most direct effects may be contributed by synaptic adhesion molecules, which are known to be involved in synaptic plasticity of sensory neurons.

We have mapped heritability to two specific types of primary sensory neurons. However, as previously mentioned, we do not explain the full heritability to musculoskeletal chronic pain in this study, since cell types other than those analyzed may also contribute to chronic pain. Furthermore, the enhancing statistical power and functional human genomic data may add resolution. Thus, more genes could contribute to chronic pain within sensory neurons as well as in cells other than primary sensory neurons, such as immune and vascular cells for headaches. Nevertheless, the finding that two neuronal types among the variety of sensory neurons carry a significant enrichment for the heritable risk to musculoskeletal pain indicates a contribution of these cell types to the pathophysiology of musculoskeletal chronic pain. This provides a rational for a deeper investigation of their participation with regards to human chronic pain. Furthermore, many drugs fall into the translational chasm between mice and humans^72^. Our atlas of somatosensory neuron gene expression in non-human primate should be important to verify putative drug targets and can also be of help in informed strategies for developing conceptually new analgesic drugs.

## Materials and Methods

### Animals

*For WaferGen (STRT-2i-seq), DRGs from two 5-year-old females (samples WG17019 and WG17020*) and one 14-year-old male (WG18008) macaques were used. Smart-Seq2 samples were prepared from five females aged 5-7 years from the same colony. All used animals were healthy Indian rhesus macaques (*Macaca mulatta*) from a colony of outbred animals housed in the Astrid Fagraeus laboratory at the Karolinska Institutet: The tissue was obtained from already euthanized animals sacrificed as a part of an unrelated study approved by the Stockholm Ethical Committee on Animal Experiments, organized under the Swedish Board of Agriculture (permit N2/15)^73,74^.

### Preparation of cell suspensions

Approximately one hour after animal termination 6-8 lumbar DRGs (3 pairs of the biggest and one pair of anterior/posterior L3-L7) were exposed, dissected out and kept in cold NMDG-cutting solution until dissociation (NMDG-CS adopted from^75^; concentrations in mM: 103 NMDG (10 N HCl to adjust pH to 7.4), 2.5 KCl, 1.2 NaH2PO4, 30 NaHCO3, 20 HEPES, 25 Glucose, 5 sodium ascorbate, 2 Thiourea, 3 sodium pyruvate, 10 N-acetyl-L-cysteine, 1 mM EDTA). Dissociation procedure was modified from^76^ and all reagents were from the same sources. All following steps were performed in cold NMDG-CS. Connective tissue and nerve excess were removed before dissociation. DRGs were cut once longitudinally (along the nerve) followed by chopping into ∼0.5 mm slices with a tissue slicer. Samples were enzymatically treated (3.9 ml papain solution (45 µ/ml, PAPL, cat.n.LS003118, Worthington) in NMDG-CS, 0.2 ml DNAse I (1 mM in NMDG-CS, cat.n.LK003172, Worthington), 0.4 ml TrypLE (cat.n.12605036, Life Technologies), 0.4 ml Collagenase/Dispase (20mg/ml, cat.n.LS004106, Worthington)) for 1 h at 37°C with triturating every 15 min through a 1 ml tip cut to a ∼5 mm opening. Then the sample was gently triturated trough 3-4 cut 1 ml tips (5 to 2 mm opening). The resulting cell suspension was then run through 100 µm cell-strainers (pluriStrainer^®^, 100 µm). The total volume of the sample (after strainer rinsing) was ∼5 ml. 1 ml of 10% Optiprep (cat.n. D1556, Sigma Aldrich) in NMDG-CS was loaded under the cell suspension solution using a gel loading tip and centrifugated at 200g for 6 min with a low break. Supernatant was discarded, the cell pellet re-suspended in NMDG-CS+B27 supplement (NMDG-CS/B27) and run through a 10 µm pluriStrainer^®^ pre-coated with NMDG-CS/B27. The strainer was rinsed twice with 5 ml of NMDG-SC/B27 applied slowly along the wall of the strainer, turned over, and the remaining big cells were then collected by back flushing the filter twice with 5 ml of filtered NMDG-CS/B27. 5 µl of CellTracker (Green CMFDA, cat.no. C7025, 50 µg in 10 µl of DMSO) was added into the resulting 10 ml of cell suspension. The cells were kept on ice for 15-20 min, centrifuged for 3 min at 100 g and re-suspended in 900 µl of 0.45 µm filtered NMDG-SC/B27/12% optiprep (optiprep was added to slow cell sinking in the solution while dispensing) and dispensed immediately into a WaferGen9600 Chip. Samples for Smart-Seq2 protocol (SS2) were dissociated in a similar way but CellTracker Orange was used instead of Green. Then samples were run through FACS (BD Influx, nozzle - 200 µm pressure - 2.5 PSI), gating events for high values in both SSC and Orange fluorescent channel and sorted into 384 plates for immediate freezing for later processing in the Smart-Seq2 pipeline.

### scRNA-seq, sequencing, and sequence alignment

WaferGen chips were processed according to the STRT-seq-2i workflow described in^77^ and FACS sorted 384-well plates were processed according to the Smart-Seq2 workflow described in^78^. The resulting WaferGen (samples WG in the text) and Smart-Seq2 (samples SS2 in the text) were sequenced on three and one lanes, respectively, on the Illumina HiSeq 2500 platform. Reads were aligned to the Macaca mulatta genome build (Mmul_10 / rheMac10, ftp://hgdownload.soe.ucsc.edu/goldenPath/rheMac10/). Since transcript annotations are incomplete on the macaque genome, and names of macaque genes differ from these of the human orthologues or putative orthologues, we used the following procedure to extend and normalize the monkey transcript annotation set with the human hg38 annotations: 1) LiftOver (http://hgdownload.soe.ucsc.edu/goldenPath/rheMac10/liftOver/) was used to align the rheMac10 and hg38 genomes. 2) A transcript from hg38 that had not even the smallest overlap with any transcript in rheMac10 was added to final transcript set. 3) All transcripts from rheMac10 were put in the final set. If either 5’ or 3’ end matched within a few bases to a human transcript, the gene name was changed to the human name. The exception to this was microRNAs (‘MIRxxx’) in hg38, where both ends were required to match for a name change to be applied. 4) In the few cases where two hg38 transcripts with the same 5’ or 3’ end had different gene names, a rheMac10 transcript with the same end was assigned a name which is a combination (‘name1/name2/…’) of the alternative names from hg38. 5) When there was a longer 5’ or 3’ extension in the liftOver model of a human transcript compared with the overlapping monkey transcript, the 5’/3’ exons of the corresponding monkey gene were adjusted accordingly. The outcome gene statistics for raw WG sample data is the following: median genes detected per neuron - 6380, total number of genes detected in at least three neurons – 22160, of them 15821 are in human gene nomenclature consortium.

### RNAScope, microscopy, image analysis and quantification

Sets of freshly dissected macaque lumbar DRG were fresh frozen in OCT over dry ice and swiftly stored at -80°C until sectioning. Cryosections were cut at 10-20µm thickness, the slides were dried at RT and then stored again at - 80°C to preserve RNA integrity. RNAScope version 2.0 using the RNAscope Fluorescent Multiplex Reagent Kit (Advanced Cell Diagnostics Inc.) was used for *in situ* hybridization. We used the staining combinations listed below for validations. TrpM8^High^: *TRPM8, SCN10A* (negative), *CBLN2* (negative); TrpM8^Low^: *TRPM8, SCN10A, CBLN2* (negative); A-LTMR: *CBLN2, TRPM8* (negative), *SCN10A* (negative); NP1: *GFRA1, GFRA2, GAL* (negative); NP2: *GFRA1, GFRA2* (negative), *GAL* (negative); C-LTMR: *GFRA2, GFRA1* (negative), *GAL* (negative); PEP1: *GAL, GFRA1* (negative), *GFRA2* (negative); NP3: *GFRA3, PVALB* (negative), *GAL* (negative); PEP2: *GFRA3, PVALB, GAL* (negative). The following probes were used to validate the clusters: Hs-SCN10A (#Cat No. 406291), Hs-TRPM8-C3 (#Cat No. 543121), Mmu-GAL-C3 (#Cat No. 830031), Hs-GFRA1-C2 (#Cat No. 435661), Hs-GFRA2 (#Cat No. 463011), Hs-CBLN2-C2 (#Cat No. 446051), Hs-GFRA3-C2 (#Cat No. 535521), Mmu-PVALB (#Cat No. 461691). The cross-reaction of human probes was predicted computationally by Advanced Cell Diagnostics Inc. and verified in macaque tissue by the authors. A Zeiss LSM800 confocal microscope was used to capture images and ImageJ (v.1.52h) for image analysis. Lumbar DRG sections from three animals (females, about 5 years old) different from those used for scRNA-sequencing were processed for quantification. Quantification of the proportion each neuronal type in the DRG and measurements of cell diameter of the different neuronal types was performed by two independent persons on the three animals (total cells analyzed = 2228 cells). Background enhancement was used to identify all neuronal profiles with a visible nucleus in each section. Quantification was performed automatically by imageJ using a probe signal threshold as positive classifier. Probe signal threshold was determined for each individual probe signal using a signal distribution plot. Different animals showed similar relative dimeter of each neuronal type (Pearson correlation coefficient = 0.98).

### scRNA-seq analysis of Macaque data

R (version 4.0.2) and Seurat (version 3.2.2) were used for the scRNA-seq analysis. Three objects were created from the individual biological STRT-2i-seq replicates. The data was normalized (*NormalizeData*) after which 2,000 most variable features were selected (*FindVariableFeatures*). To mitigate batch effects between replicates we used Seurat’s integrated analysis approach that transforms datasets into a shared space using pairwise correspondences (or ‘anchors’)^40^. Anchors were first defined using *FindIntegrationAnchors* (dims = 1:40) and the data was then integrated (*IntegrateData*) and scaled (*ScaleData*), followed by principal component analysis (PCA) (*RunPCA*, npcs = 100). For clustering, the following parameters were used: *RunUMAP*, reduction = pca, dims = 1:20; FindNeighbors, reduction = pca, dims = 1:20; *FindClusters*, resolution = 1. The clusters showing minimal expression of a neuronal markers (*SNAP25, RBFOX3*) and/or low number of detected genes were removed. The following round of clustering (resolution = 3) produced a total 29 clusters of which three expressing markers of injured cells (e.g. *ATF3*) satellite glia (e.g. *APOE*) or other non-neuronal cells (e.g. *EMCN* [endothelial cells]) were removed. The cells were then reclustered (resolution = 2) and remaining cells with ambiguous identities were removed. After this, a residual signal coming from satellite-glia marker genes was scored for each cluster (*AddModuleScore*; features = “SPARC”,”PMP2”, “APOE”, “SPARCL1”, “PLP1”, “ABCA8”, “S100A1”, “LPL”, “IFITM4P”, name = “scg_score”). The cells were then clustered for the final time (dims = 1:30, resolution = 2) and the satellite-glia signal was regressed out (*ScaleData*; vars.to.regress = “sgc_score1). After this, highly similar clusters without clearly distinguishable markers were merged to produce the final nine clusters. A dendrogram was then built (*BuildClusterTree*) using dimensions 1:30. For Smart-seq2, the individual datasets for each animal were initially integrated and clustered similarly to the STRT-2i-seq data. After the first round, clusters showing a non-neuronal identity and/or low levels of detected genes were removed. After this, the cells were cluster a second time (dims = 1:20, resolution = 1.5). To assign identities to the clusters we used label transfer between the STRT-2i-seq and Smart-seq2 datasets (*FindTransferAnchors*; reference = STRT-2i, query = Smart-seq2, dims = 1:30; *TransferData*; anchorset = anchors, refdata = usoskin_id, dims = 1:30)). Wilcoxon rank sum test was used to identify cluster-enriched genes, reporting adjusted p-values based on Bonferroni’s correction (false discovery rate, FDR). Genes detected in > 25% of cells in the cluster with FDR < 0.05 and > 0.25 average log2-fold change were considered as marker genes for the cluster.

### Label transfer between mouse DRG datasets

To visualize mouse DRG scRNA-seq datasets as UMAPs, the individual datasets from^7,9^ were clustered with Seurat (*ScaleData*, vars.to.regress = c(“orig.ident”, “nCount_RNA”); *RunPCA*, npcs = 100; *RunUMAP*, dims = 1:20; *FindNeighbors*, dims = 1:20; *FindClusters*, resolution = 0.1) but the cell identities were set as original identities from the publications. Label transfers between the datasets were then performed using the different combinations of reference and query datasets and nomenclatures (*FindTransferAnchors*; reference = reference_dataset, query = query_dataset, dims = 1:30; *TransferData*; anchorset = anchors, refdata = used_nomenclature, dims = 1:30).

### Conos analysis

For Conos^41^ WG macaque datasets were integrated using CCA space ($ buildGraph(k=15, k.self=5, space=‘CCA’, ncomps=30, n.odgenes=2000, snn=F, snn.k = 50), followed by UMAP embedding ($ embedGraph(method=“UMAP”, min.dist=0.1, spread=20, n.cores=4, min.prob.lower=1e-3). For macaque (WG) and mouse (Zeisel et al.) datasets co-integration different parameters were used: $ buildGraph(k=15, k.self=5, space=‘CCA’, ncomps=30, n.odgenes=2000, snn=T,snn.k = 15), followed by largeVis embedding: $ embedGraph(alpha=0.1, sgd_batched=1e6, seed = 1).

### Mouse DRG scRNA-seq data

Mouse DRG data was downloaded from (http://loom.linnarssonlab.org/clone/Mousebrain.org.level6/L6_Peripheral_sensory_neurons.loom) and GEO (GSE139088) and the cluster annotation was modified to conform to established nomenclature^8^. Mouse gene names were switched to corresponding human orthologs using biomaRt (v2.42).

### Scoring analysis of cell-type identity

For this analysis, our goal was to score the probabilistic cell identity of each cell relative to the defined cell types at the transcriptional level^79,80^. We built a vanilla neural network model for classification tasks in PyTorch framework with CUDA support for GPU computation, and trained the model to learn the general prototypes of defined cell types. To train the model, we obtained the over-dispersed genes by estimating the mean and coefficients of variation. The over-dispersed genes were further ranked by two heuristics for cell-type specificity of both fold-change and enrichment score-change^9^. The cross-species alignment was performed as described in^80^. Subsequently, the ranked marker genes of defined cell-types were log-transformed and scaled by Minmax normalization, and then used for the neural network model. The neural network model contains an input layer with the number of neuron nodes similar to the number of marker genes, a hidden layer with the number of neuron nodes similar to 20% of marker gene numbers, and an output layer with the number of neuron nodes similar to the number of defined cell types. Linear regression was performed between each layer, and the 30% threshold for dropouts was set to reduce the overfitting. Rectified linear units were used as the activation function of the hidden layer, and Softmax was used for the output layer to evaluate the probabilities. Nesterov Momentum was used as stochastic gradient descent (SGD) optimizer. To choose the adequate regularization strength, the classifier accuracy and the loss value were inspected against epoch numbers. The classifier accuracy was estimated by a k-fold cross-validation, of which the dataset was randomly split (k=3). The learning rate, epoch number, and momentum were chosen corresponding to the maximum point of learning curve reaching the accuracy plateaus. The ready model was used to predict the probabilities of each cell belonging to each trained reference cell-types. The permutation test of dataset was applied to qualify the significance of the prediction, and the p-values were calculated by FDR. Data were visualized using the radar plot^81^ consisting of a sequence of equiangular polygon spokes with the distal vertex representing each trained reference cell-type. The distance between the polygon center and each vertex of the polygon represents the relative probabilities of each trained reference cell assigning to the defined reference cell-types and the position of each predicting cell was calculated as a linear combination of the probabilities against all reference cell types, and then visualized as the relative position to all vertices of the polygon.

### Comparison of transcriptional signatures between species

Individual Seurat objects were formed for each species from the two mouse datasets and from the two macaque datasets (Seurat function *merge*). Within each object, the genes expressed in each cluster were first filtered (*FindMarkers*; only.pos = TRUE, logfc.threshold = 0). Then, genes found specifically in one species (*setdiff*; species1$ genes, species2$ genes) or in both (intersect; species1$ genes, species2$ genes) were listed. A log2Fc threshold of 0.25 was used for finding genes that were expressed above baseline for a corresponding cluster in both species. A similar threshold was used when defining species unique cluster markers.

### SCENIC

SCENIC^43^ was used to infer active transcription factors and their target genes. SCENIC was performed following the author’s vignette: https://github.com/aertslab/SCENIC/blob/master/vignettes/SCENIC_Running.Rmd. First, gene sets co-expressed with transcription factors were identified and modules kept if the transcription factor motif was enriched among its targets. Then, target genes without direct motif support were removed. Finally, regulon activity was scored and binarized to determine whether the genes in each regulon are enriched in each cell.

### Computational genomics screen for gene families

For the supervised computational screen using Metaneighbor^82^, a set of 1500+ gene families was downloaded from the HGNC website (https://www.genenames.org/download/custom/). In brief, this is a machine-learning based analysis scores the performance to correctly connect cells of known corresponding identity based on the similarity of their transcriptional profiles across a given set of genes^42^. The analysis was run following the vignette provided by the authors (https://github.com/maggiecrow/MetaNeighbor). To improve readability, the lists of highest scoring families were curated for redundancy for the figures (Extended Data Fig. 8), showing only the highest scoring family from a group of highly similar families. The full tables can be found in Extended Data Tables 5, 6 and 7.

### Cell-type specific partitioned heritability

We followed the protocol established by^32^. Three steps of analysis were required to address our hypothesis. The first step consisted in identifying the top 10% genes most associated with a macaque DRG cluster cell type. For this, we extracted from the expression matrix the raw counts of genes in each single-cell sample. Sample size factors were estimated using DESeq2’s estimateSizeFactors^83^. Then, for each gene, we regressed gene expression of all samples to cluster membership (yes=1, no=0), using size factors and animal’s sex as co-variables. We then sorted the genes by decreasing regression’s test statistics and retained the top 10% (n=1,322) most associated genes to a cluster. The second step consisted in creating cluster-specific annotations following instructions given at URL https://github.com/bulik/ldsc/wiki/LD-Score-Estimation-Tutorial, under the “Partitioned LD Scores” section. The control track was made up of the same genes used in Franke’s control dataset^84^ by Finucane and colleagues^32^. The third and final step consisted of screening various pain-related summary GWAS from the UK Biobank against these annotations to uncover cluster-specific contributions to partitioned heritability. Statistical significance was established at up to the 20% false discovery rate (FDR) level on a per-cluster basis, i.e. the genetic contribution for each pain site was tested in each cluster. We chose FDR instead of Bonferroni as epidemiological data establish co-morbidity of pain sites report. Meta-analyses of partitioned heritability coefficients were performed using the inverse standard error approach suggested in METAL^85^.

### Cluster and GWAS enrichment

To uncover genes simultaneously strongly associated with cluster specificity and chronic pain phenotypes, we estimated the area spanned by a given gene using a scatter plot in which both cell type and GWAS enrichments were tracked. Gene-level test statistics were obtained from summary chronic pain GWAS using MAGMA^86^. We retained genes for which cell type specificity was positive, indicating an increased gene expression in the cell type of interest, with enrichment increasing with cell type specificity. P-value for a gene’s area was estimated from a fit of the density of genes to a Bessel function (appropriate to model a product of two normal distributions), then integrating from minus infinity to minus the absolute value of the area of the gene. The Bessel estimator was such that integrating over from minus infinity to plus infinity yielded a value of 1 (the estimator was normalized). All genes up to FDR 20% were retained for pathway analyses. Pathway analyses were conducted using the hypergeometrical test for over-representation^87^, with pathways sourced from Gene Ontology’s biological processes^88^, obtained December 2019 from URL http://download.baderlab.org/EM_Genesets/. We tested pathways that featured a number of genes between 10 and 1000.

### UK Biobank

The UK Biobank is a large genetic study comprised of half a million participants aged between 40 and 69 years old^47,48^. The first round of standard genotyping quality control was performed by the UK Biobank consortium and is fully documented on their web portal (http://biobank.ctsu.ox.ac.uk/crystal/refer.cgi?id=155580). Samples were discarded based on failed genotyping QC (heterozygosity rate, genotyping rate, etc.), genetic vs declared sex mismatch, voluntary retraction, and of non-“white British” ancestry. Genome-wide association study was performed using the full release 500K cohort version. BOLT-LMM version 2.3 performed association tests, with consideration of kinship among participants^89^. Age, age squared, sex, dummy-coded recruitment sites, genotyping platform, and the top 40 principal components were used as co-variables. The phenotype was defined as chronic pain at a body site (e.g. back). Cases were defined as those who answered “yes” at the touchscreen question “Have you had back pains for more than 3 months?” and that had answered “yes” at the question “In the last month have you experienced any of the following that interfered with your usual activities?” (field 6159). Control individuals were those who answered “None of the above” at field 6159 (n=163,825). A total of eight pain sites were available: headaches (UK Biobank field 3799; 36,381 cases), facial (field 4067; 3,495 cases), neck and shoulder (field 3404; 63,933 cases), stomach and abdominal (field 3741; 18,749 cases), back (field 3571; 70,634 cases), hip (field 3414; 35,634 cases), knee (field 3773; 68,237 cases), and widespread (field 2956; 5259 cases). To control for overlap of chronic pain sites in individuals, we also performed additional GWASes for pain sites in which cases who reported a particular additional pain site we removed, one site at a time, from the analysis. A total of 8*8 – 8 = 56 such GWASes were performed. Post-GWAS SNP quality control included: (1) be part of the Haplotype Reference Consortium^90^, (2) minor allele frequency > 0.1%, (3) Hardy-Weinberg P-value > 10^−12^, (4) INFO score > 0.9. In all, this yielded a total of about 8.238 million SNPs per GWAS. The minimum effective count of minor alleles was: 0.1% * 0.9 * 163,835 * 2 ∼ 295. This work was done under the UK biobank application number 20802.

### Data availability

The raw and processed datasets reported in this study have been deposited in the Gene Expression Omnibus (*GEO) under the accession* GSE148238.

### Code availability

Any custom code used in the analyses is *available* from the *authors* upon *a reasonable request*.

## Acknowledgements

We thank Martin Häring for advice on enzymatic treatment/dissociation strategies and Ming-Dong Zhang for advice on dissection. We also thank Gunilla B. Karlsson Hedenstam for donating euthenized animals, Sten Linnarsson for computational support and Eneritz Agirre for help with the Shiny-app and running SCENIC. We acknowledge the Eukaryotic Single Cell Genomics (ESCG) Facility and the Mass Cytometry National Facility at the Science for Life Laboratory, Sweden, for the scRNA-seq and for the pilot sample FACS, respectively. This work was supported by the Swedish Medical Research Council, Knut and Alice Wallenbergs Foundation (Wallenberg Scholar and Wallenberg project grant), SFO grant (StratNeuro), Wellcome Trust (Pain Consortium), European Research Council advanced grant (PainCells 740491), and Karolinska Institutet (to P.E.), EMBO fellowship to MF, and by the Canadian Excellence Research Chairs (CERC) Program (to LD).

## Author contributions

Design of experiments: J.K, D.U and PE; Computational analysis: J.K, D.U., P.K and Y.H; LiftOver, data quality assessment and gene annotation: N.B, P.K and P.L.; Design of RNAScope Experiments: J.K., M.F. and D.U.; RNAScope Experiments: D.L., D.U.; Single-cell suspensions: D.U.; Provision of primate tissues: K.L and; Dissections: D.U., M.S. and B.E.; Human genetic studies: M.P. performed bioinformatics analyses and S.K. and L.D. interpreted the bioinformatics results. Writing – Review & Editing: J.K. and P.E., with input from all authors; and Supervision and Funding: P.E.

## Competing interests

The authors declare no competing interests.

## Extended Data Figs

**Extended Data Fig. 1.**
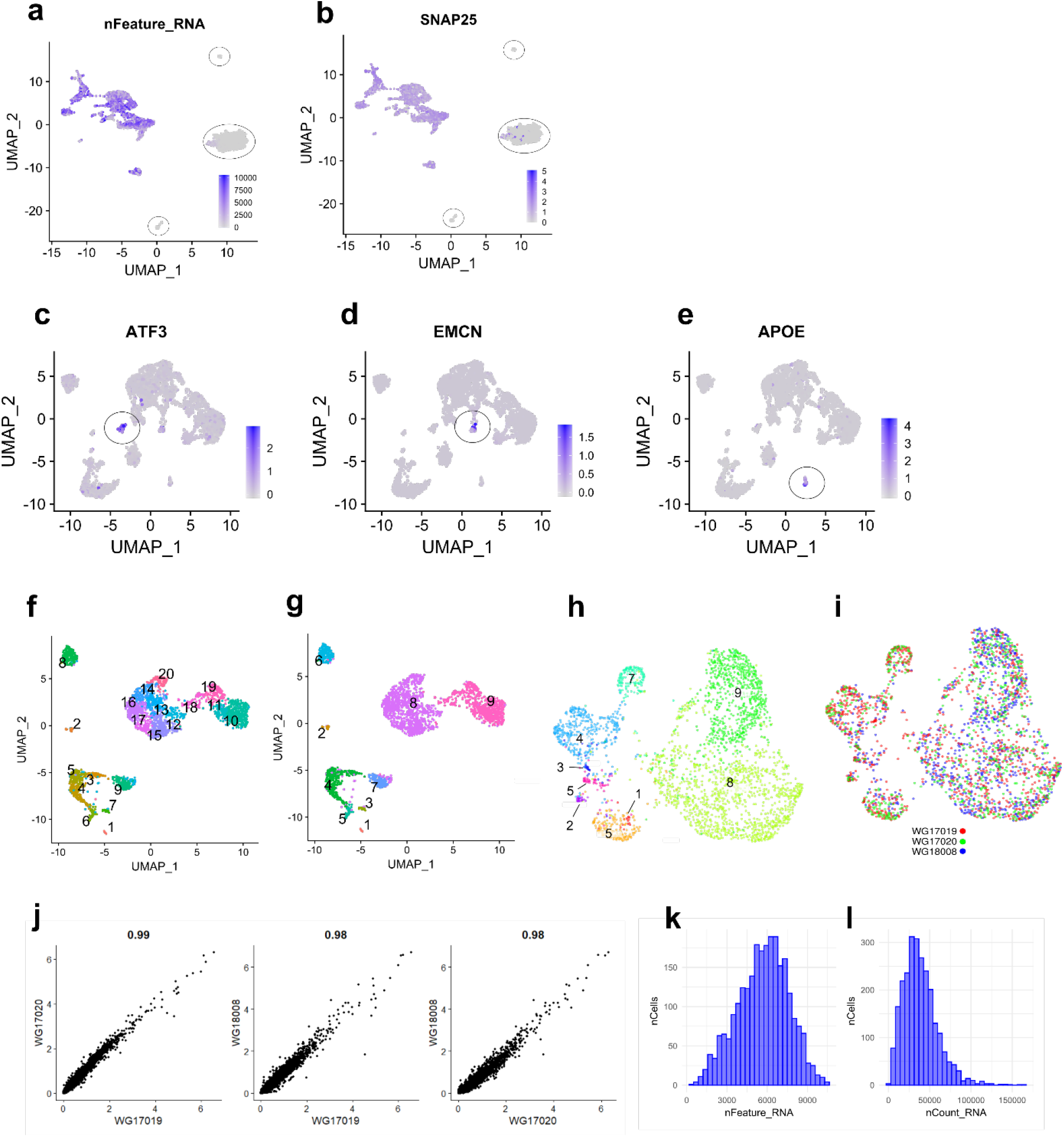
Clustering of macaque STRT-2i-seq data. (a, b) UMAPs showing removed clusters (circled) based on low gene detection (a) and lack of neuronal identity (b). (c-e) UMAPs showing removed clusters (circled) based on injury (c) non-neuronal (d) and satellite-glia (e) markers. Genes shown are representative of several marker genes. (f) UMAP showing the final 2,518 neurons after over-clustering. (g) UMAP showing the final nine neuron clusters after merging highly similar types. (h-i) UMAPs showing clustering results using an alternative method (Conos) (h) Conos clusters with numbering from original analysis showing near perfect match (i) UMAP showing the equal distribution of cells from different individuals among the clusters. (j) Scatterplots showing linear correlation of inter-individual gene expression. The Pearson correlation coefficient is shown above each plot. (k-l) Histograms showing distribution of (k) detected genes and (l) UMIs in the STRT-2i-seq data.

**Extended Data Fig. 2.**
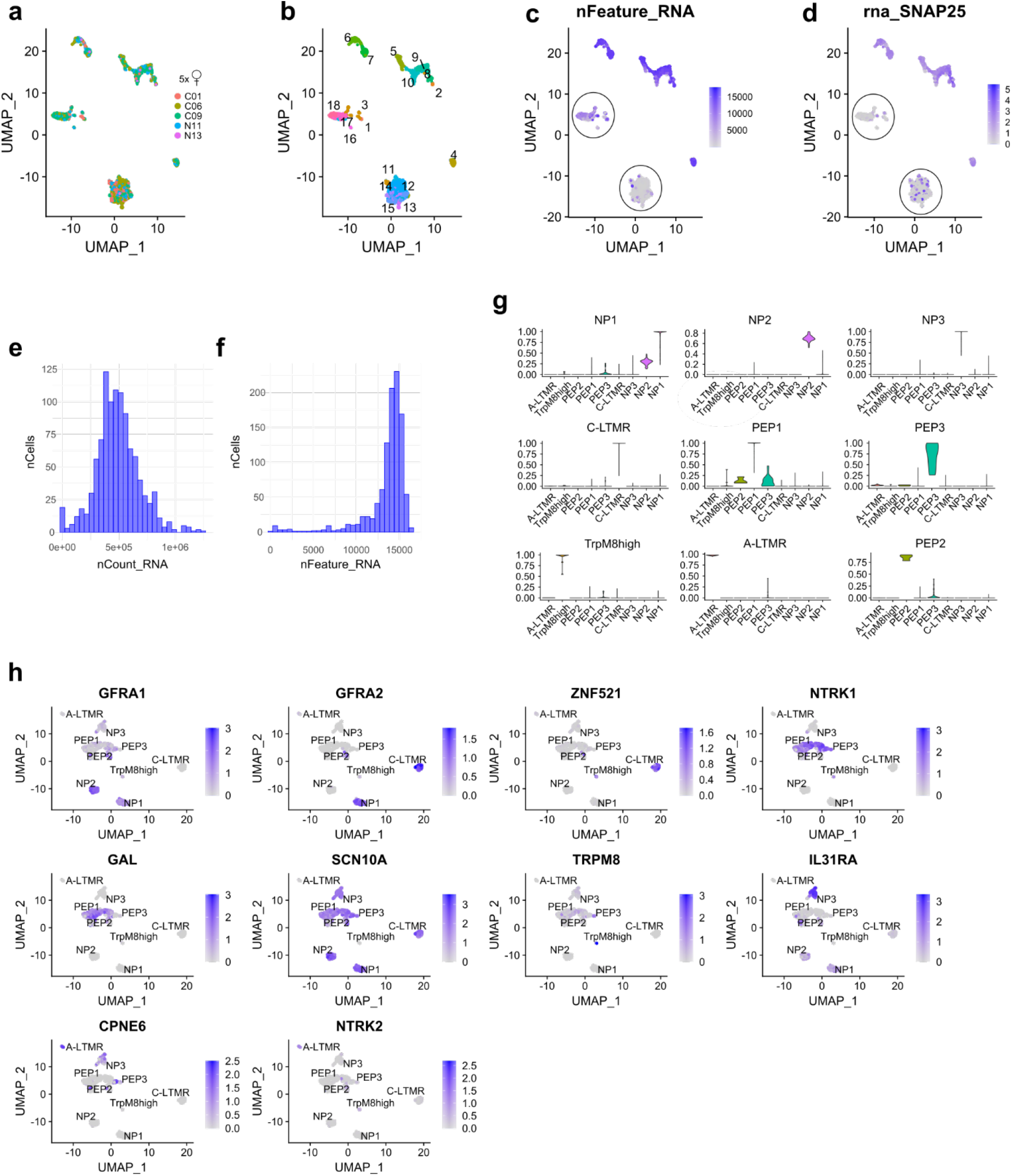
Analysis of macaque Smart-seq2 data. (a) UMAP distribution of cells originating from the individual animals after clustering. (b) UMAP showing numeric labels after primary clustering. (c, d) UMAPs showing removed clusters based on (c) low gene detection and (d) lack of neuronal markers (circled clusters). (e, f) Histograms showing the (e) transcript count and (f) detected gene count distribution in the Smart-seq2 data. (g) Violin plots showing the prediction scores from the label transfer between the STRT-2i-seq and Smart-seq2 data. (h) UMAPs showing mouse canonical marker gene expression in the Smart-seq2 macaque clusters after label transfer.

**Extended Data Fig. 3.**
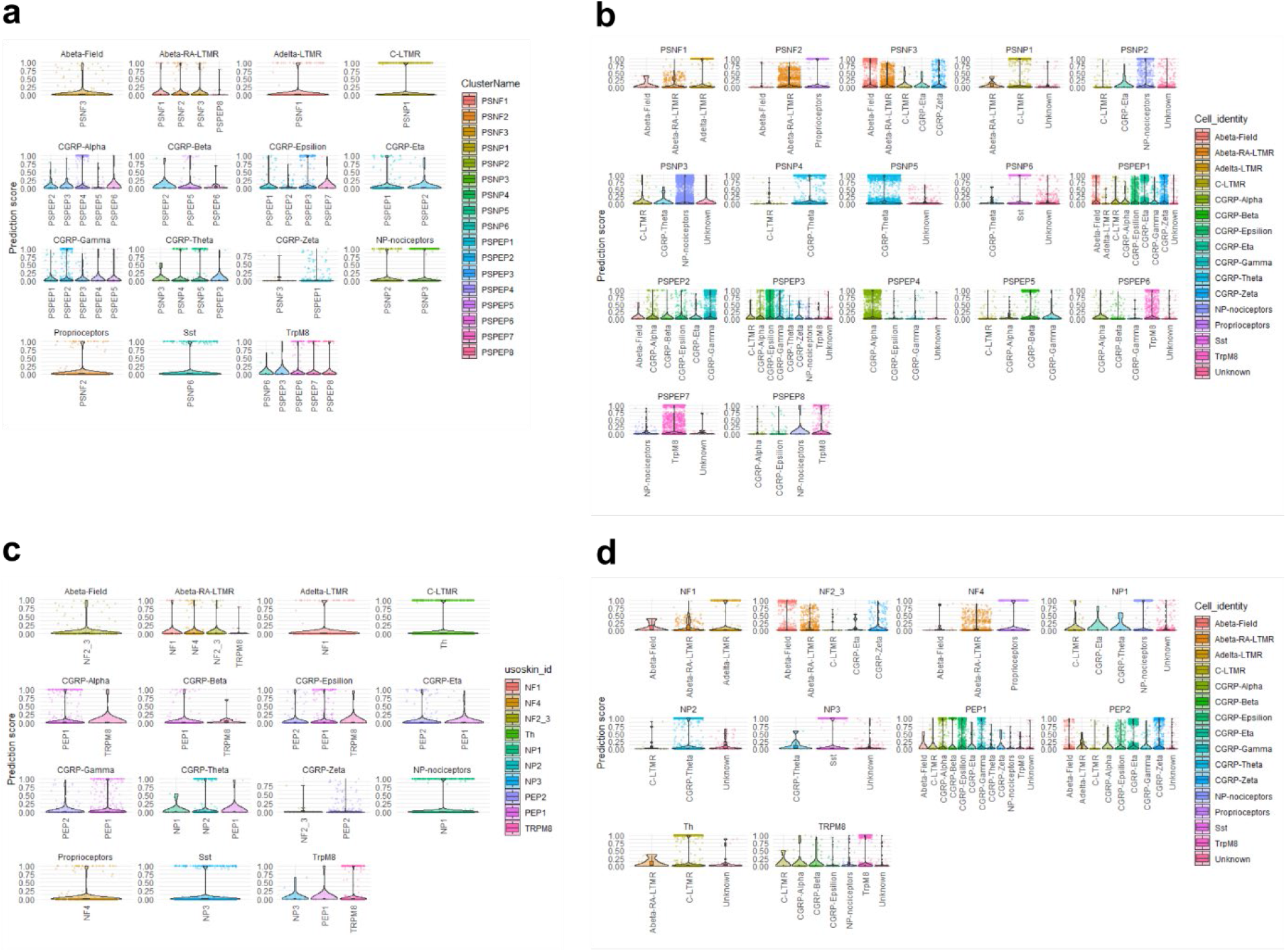
Prediction scores from label transfer between Zeisel et al. and Sharma et al. data. (a, b) Violin plots showing prediction scores from the label transfer from (a) Sharma to Zeisel and (b) Zeisel to Sharma. (c, d) Violin plots showing prediction scores from the label transfer from (a) Sharma to Zeisel and (b) Zeisel to Sharma when using Usoskin.

**Extended Data Fig. 4.**
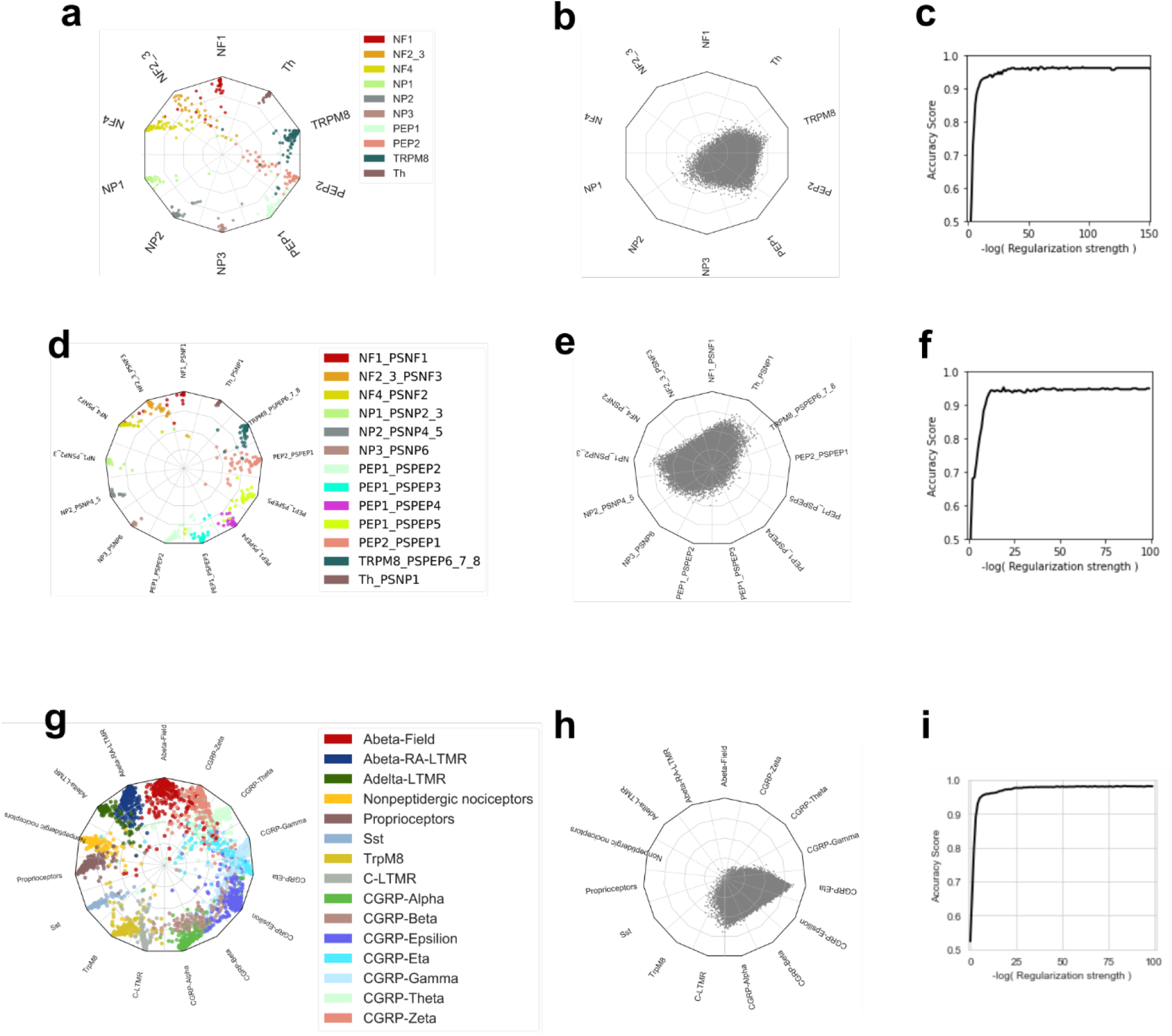
Scoring modules for prediction of sensory neuron types. (a) A radar-plot showing cell-type fractions of the reference mouse DRG neurons (1/3 of cells) from Zeisel et al.^9^ from the neural-network scoring module as described in Materials and Methods. Each dot represents one cell, and the color coding is based on unique cell clusters. (b) Negative control obtained from the same module after random permutation of the features in the training dataset and visualized in the radar plot. (c) Accuracy score for the learning module. (d-e) Scoring module from Zeisel et al. data using original Zeisel nomenclature. (g-i) Scoring module from Sharma et al. data using original Sharma nomenclature.

**Extended Data Fig. 5.**
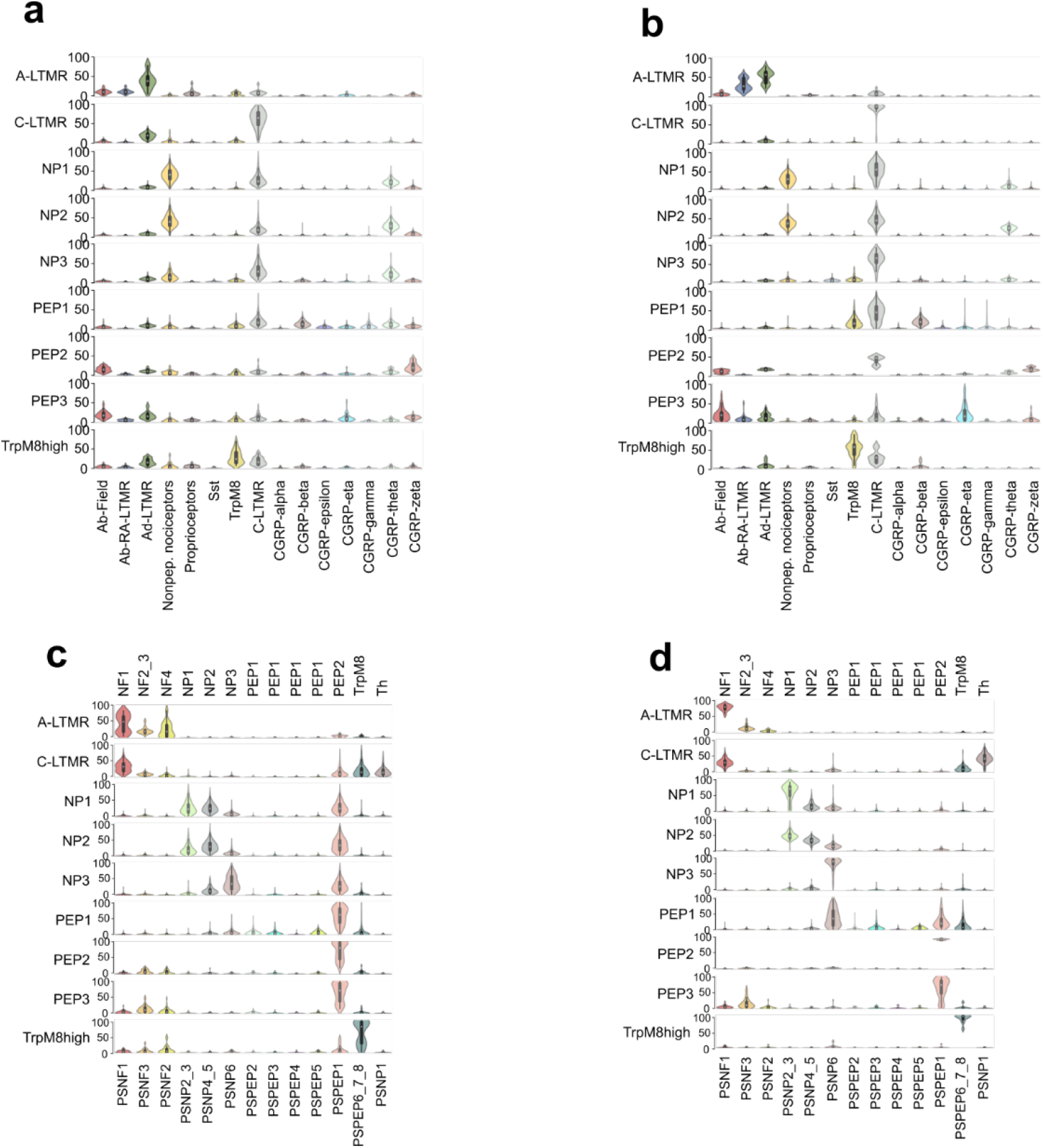
Prediction of macaque DRG neuron types from mouse data. Violin plots showing prediction probability scores between (a) macaque types (STRT-2i-seq) and Sharma et al. mouse types, (b) macaque types (Smart-seq2) and Sharma et al. mouse types, (c) macaque types (STRT-2i-seq) and Zeisel et al. mouse types, (d) macaque types (Smart-seq2) and Zeisel et al. mouse types. (c) and (d) have Zeisel and Usoskin nomenclatures on bottom and top, respectively.

**Extended Data Fig. 6.**
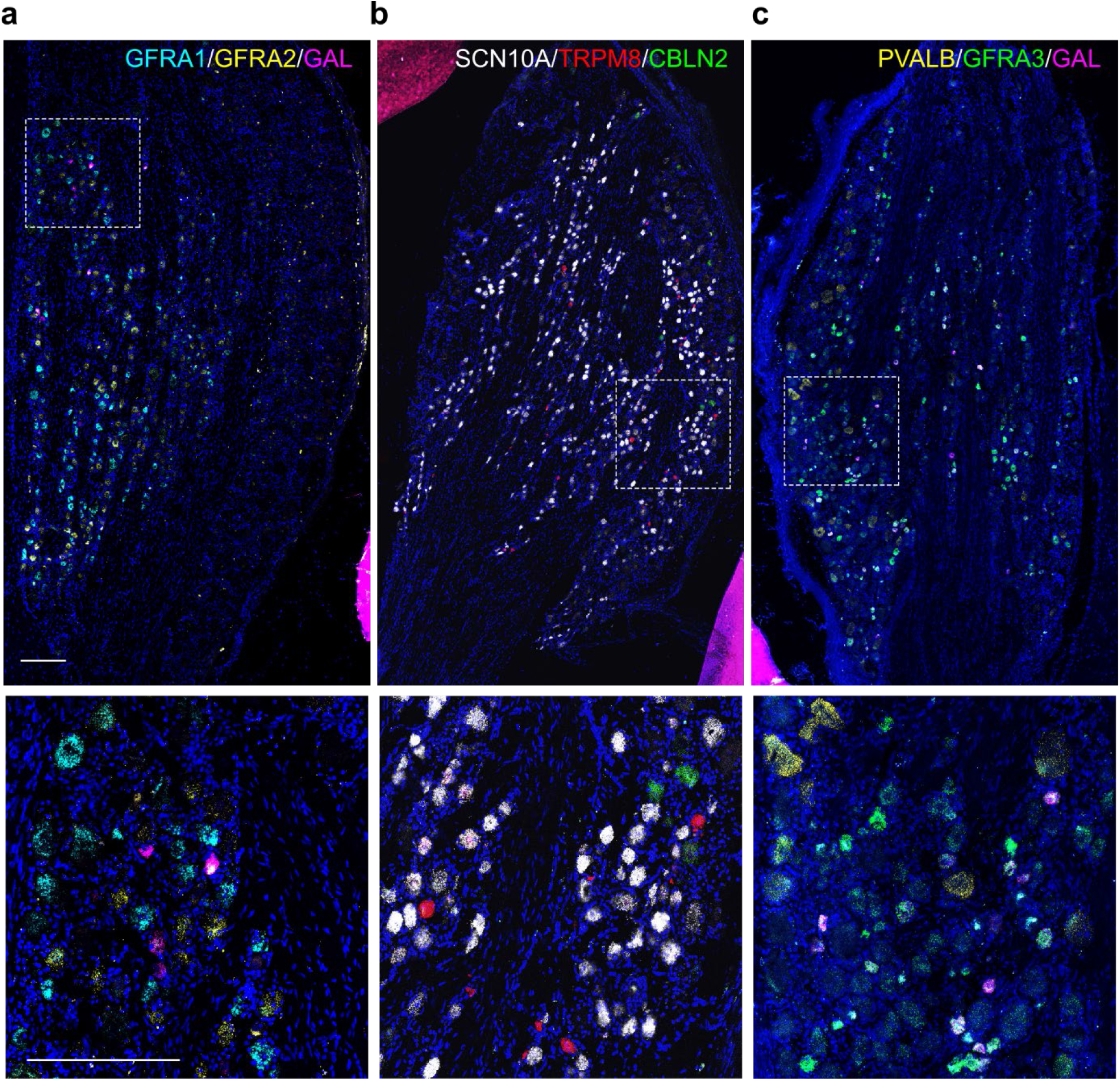
Validation of macaque clusters by triple in situ hybridization. (a-c) Images of entire Macaque DRG sections after performing triple in situ hybridization using probes specific for either (a) GFRA1 (light blue), GFRA2 (yellow), and GAL (purple), (b) SCN10A (white), TRPM8 (red), and CBLN2 (green), or (c) PVALB (yellow), GFRA3 (green), and GAL (purple). The white hatched box in the top image is shown at higher magnification in the bottom panel. Scale bar = 250 μm.

**Extended Data Fig. 7.**
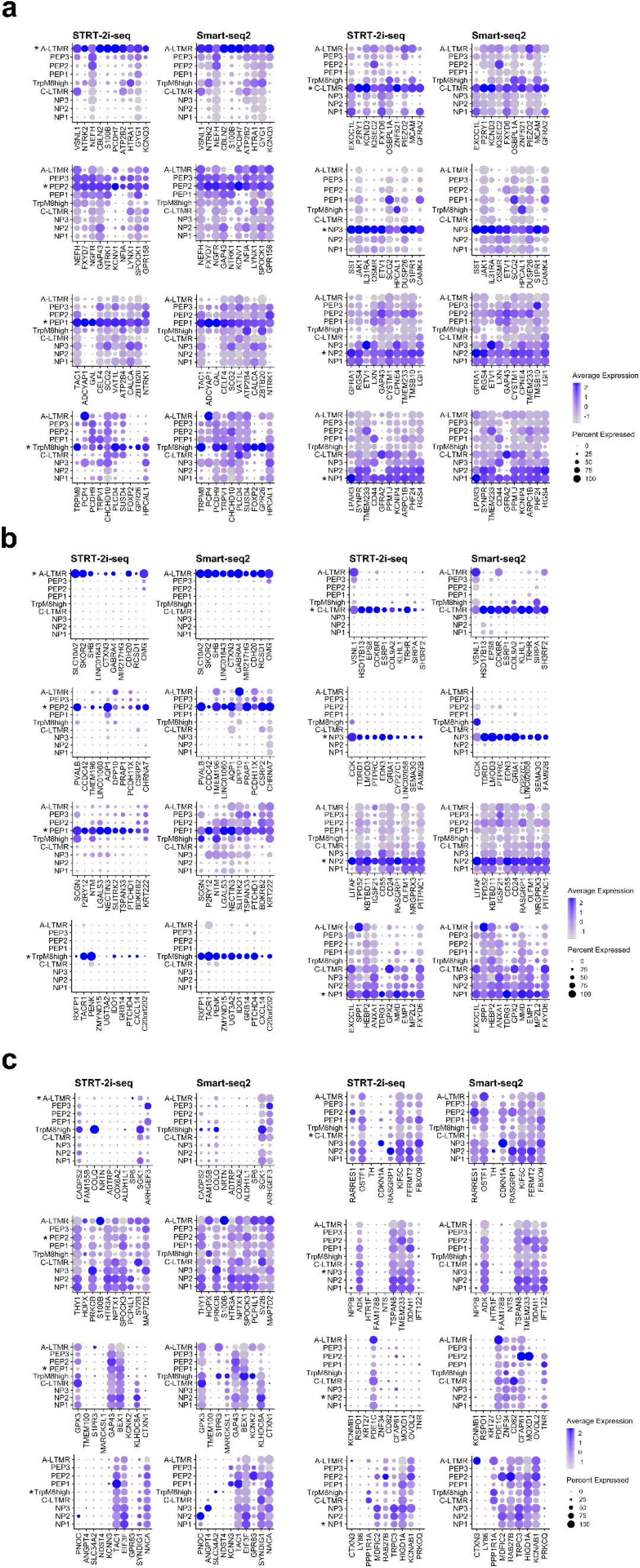
Side-to-side gene expression comparison between STRT-2i-seq and Smart-seq2 data. (a) Top genes shared between corresponding cell types of macaque and mouse plotted in both macaque datasets. (b) Top macaque enriched genes plotted in both macaque datasets. (c) Top mouse enriched genes plotted in both macaque datasets. Cell type to compare is marked with an asterisk in each panel.

**Extended Data Fig. 8.**
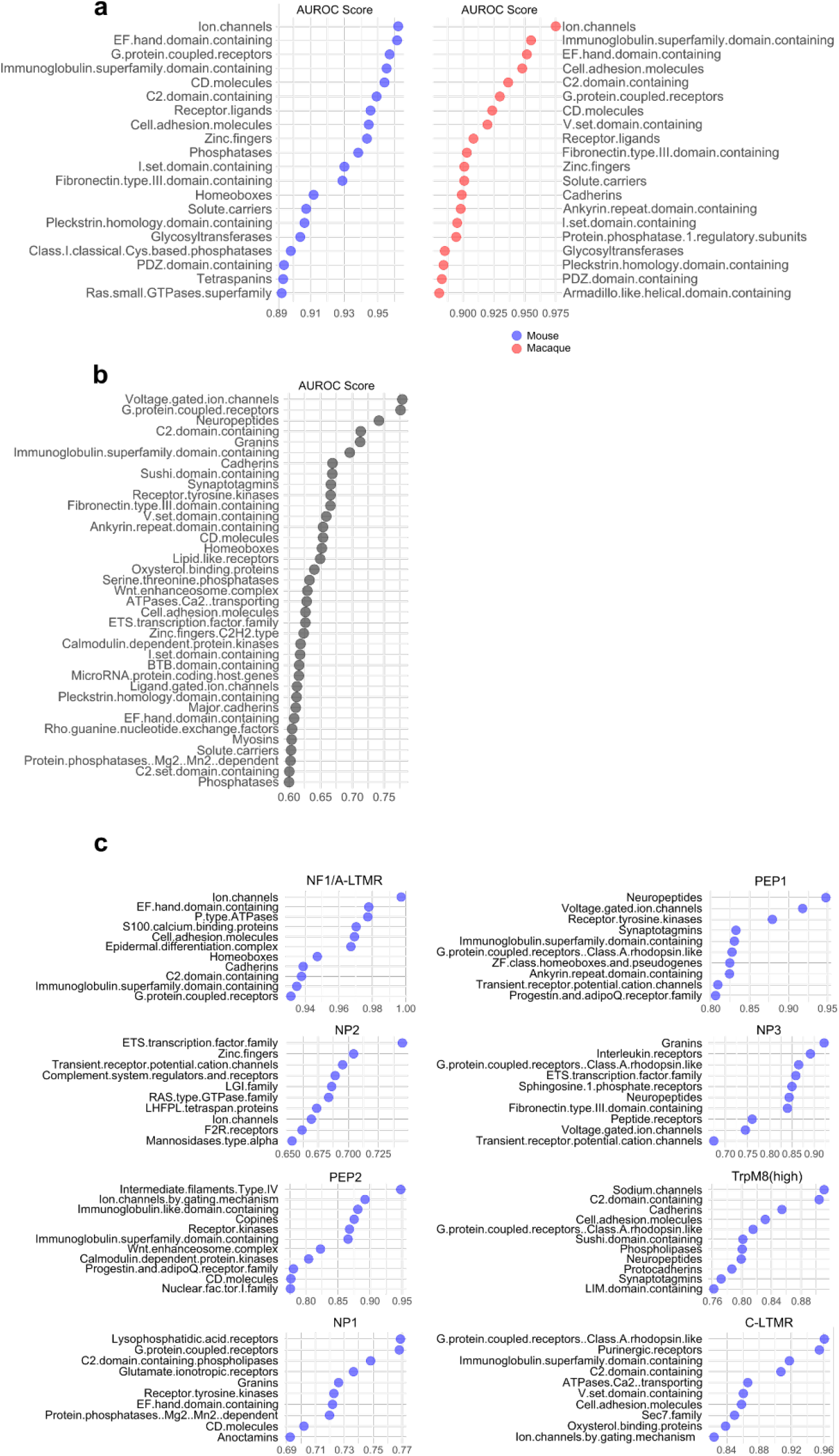
Computational screen of gene families that define DRG cell types. (a) Gene families showing high performance in correctly assigning cell types in both macaque and mouse. x-axis shows the average AUROC score over all cell types. (b) Gene families showing the highest performance in correctly assigning cell types between corresponding cell types of macaque and mouse. x-axis shows the average AUROC score over all cell types. (c) Gene families showing the highest performance in correctly assigning cell types between specific corresponding cell types of macaque and mouse. Note that the lists in the figure have been curated to remove highly redundant families. The full lists are available in Extended Data Tables 5 through 7.

**Extended Data Fig. 9.**
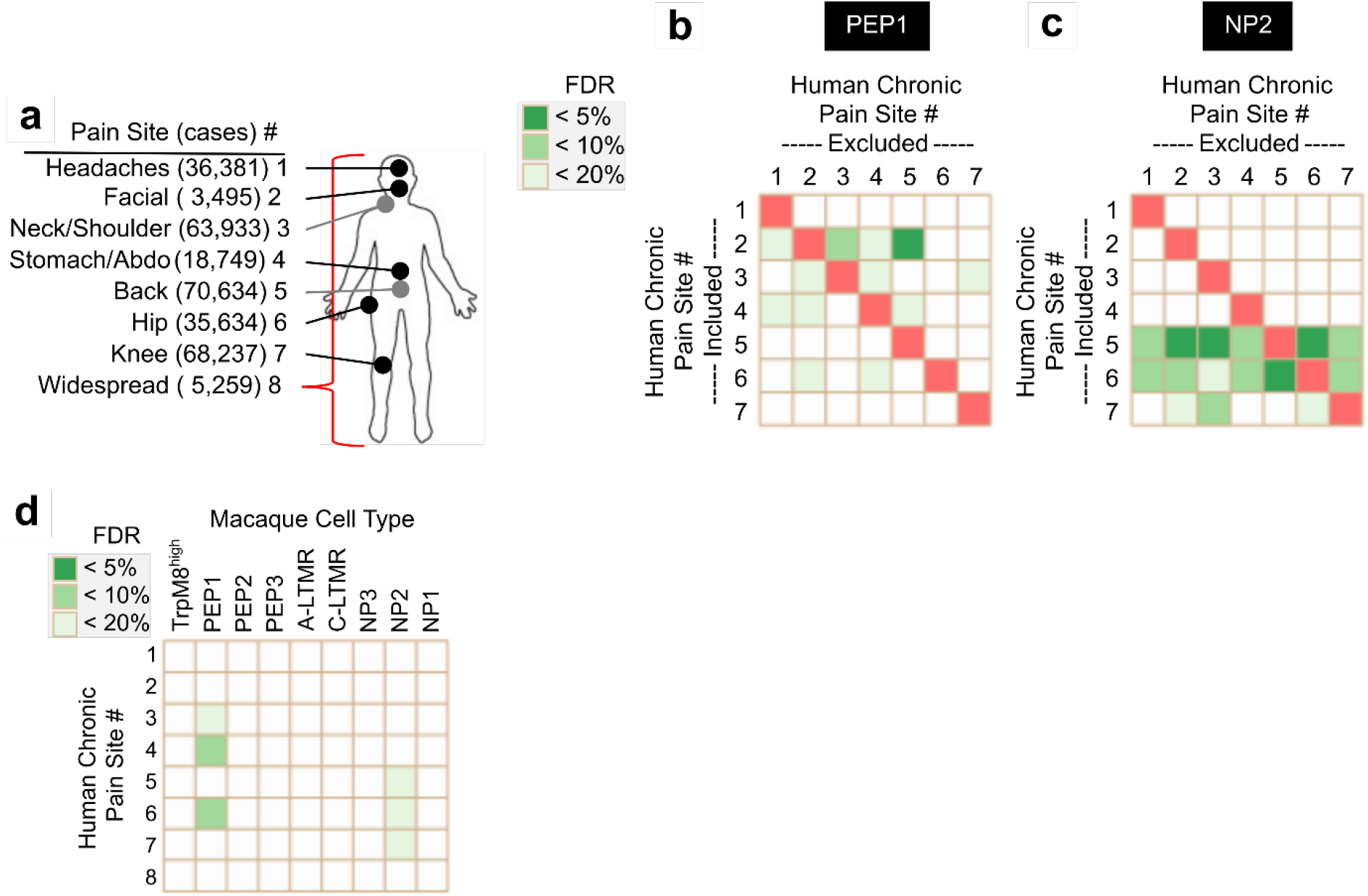
Partitioned heritability analysis in the Smart-seq2 data and leave-one-out analysis of partitioned heritability in the STRT-2i-seq data. (a) UK Biobank chronic pain sites mapped to the human body. Number of chronic case participants shown in parentheses. (b, c) Heatmaps of FDR-corrected p-values for enrichment in partitioned heritability of a macaque DRG cell type to each human chronic pain site. For a particular chronic pain site GWAS (row), the cases reporting other chronic pain site (column) were removed from the that GWAS. Widespread pain cases do not overlap with any other cases for the other sites (mutually exclusive questionnaire choices). (d) Heatmap of FDR-corrected p-values for enrichment in partitioned heritability of each macaque DRG cell type to each human chronic pain site in the Smart-seq2 data.

## Notes

### Competing Interest Statement

The authors have declared no competing interest.

